# Specialized Computations for Generalized World Modelling in Medial Prefrontal Cortex

**DOI:** 10.64898/2026.04.15.718798

**Authors:** Fahd Yazin, Gargi Majumdar, Christopher G. Lucas, Neil R. Bramley, Paul Hoffman

## Abstract

The medial prefrontal cortex is central to learning flexible internal models across diverse domains, yet the functional specialization enabling this remains unknown. We tested whether medial prefrontal specialization is representational (encoding domain-specific features) or computational (implementing domain-general computations). During fMRI, participants learned probabilistic features of virtual environments representing spatial, social, and sequential domain knowledge. Although each domain used different features, they shared the same feature-to-latent state mapping, matching their computational demands. The medial PFC showed no domain-specific feature representations. Instead, its neural patterns revealed a triad of specialized yet domain-invariant computations. Ventromedial PFC patterns reflected probabilistic inference, abstracting hidden probability distributions from observations and tracking trial-wise posterior task state changes within a low-dimensional latent space. Anteromedial PFC organized task states along orthogonal axes, tracking directional shifts within states and switches between different states, suggesting a global task coordinate system. Dorsomedial PFC patterns represented task dynamics, using predictive surprise to monitor validity of the current internal model and switch task policies. These results suggest a principled architecture in medial PFC where three general-purpose computations jointly enable learning world-models across diverse environments.

## Introduction

Our experience of the world is often incomplete and contradictory. Yet from sparse, noisy and varying perceptual inputs, the brain is able to extract statistical regularities^1^. Learning these regularities enables us to adaptively construct internal models of the world that distil and generalize key features of our experience. For example, when lower-order visual statistics are insufficient for optimal learning, higher-order regularities are automatically computed^2^. Such efficiency suggests we use a repertoire of inductive biases, optimized for rapid learning from limited data. Many of these biases appear tuned to specific perceptual domains^3^. Object understanding benefits from continuity and permanence priors that emerge early in life, even before extensive visual experience^4^. Social perception, similarly, is aided by biases for recognising faces^5^ and biological motion^6^. It is unknown, however, whether inductive biases also shape the acquisition of abstract internal models. The midline prefrontal cortex (PFC ; throughout the paper, we use the abbreviation PFC to refer to medial (and not lateral) prefrontal regions) is known to learn generalizable internal representations^7– 9^, and its functional heterogeneity suggests it may contain a range of biases that aid learning. Here we investigate whether and how the midline PFC expresses computational preferences when humans learn latent properties across three disparate domains (social, spatial and sequential)^1,10^.

Midline PFC forms part of the Default-Mode Network (DMN), which is generally implicated in internally-driven cognition and the formation of mental models^11–13^. It has various subregions which are sensitive to distinct environmental variables (Fig 1e). Ventromedial PFC (vmPFC) is involved in representing the properties of specific spatiotemporal contexts^14–16^. Anteromedial PFC (amPFC) is implicated in social cognition tasks such as inference of others’ mental states^17–20^ and inference of positions within a social hierarchy^19^. And finally, dorsomedial PFC (dmPFC) is implicated in action observation and formulating action plans^21,22^. Indeed, in recent work, we observed a functional dissociation of this form during naturalistic movie-viewing and narrative listening^23^. Specifically, belief updates about contextual states, social references, and action transitions in the movie or narrative were preferentially associated with activity peaks in vmPFC, amPFC, and dmPFC, respectively. The most common interpretation of such findings is that they indicate functional specialisation within midline PFC for particular domains of experience. However, an alternative possibility is that midline PFC regions vary in their inductive biases, such that functional specialisation is driven by variation in computational properties across the region, not by sensitivity to different domains per se. On this view, apparent domain-specific activation arises because understanding in different domains tends to rely on different types of computation. In the next section, we consider the functional specializations that may be relevant in each domain. We then describe how we tested this account in a learning task, by varying domain while holding computational demands constant.

**Fig 1:**
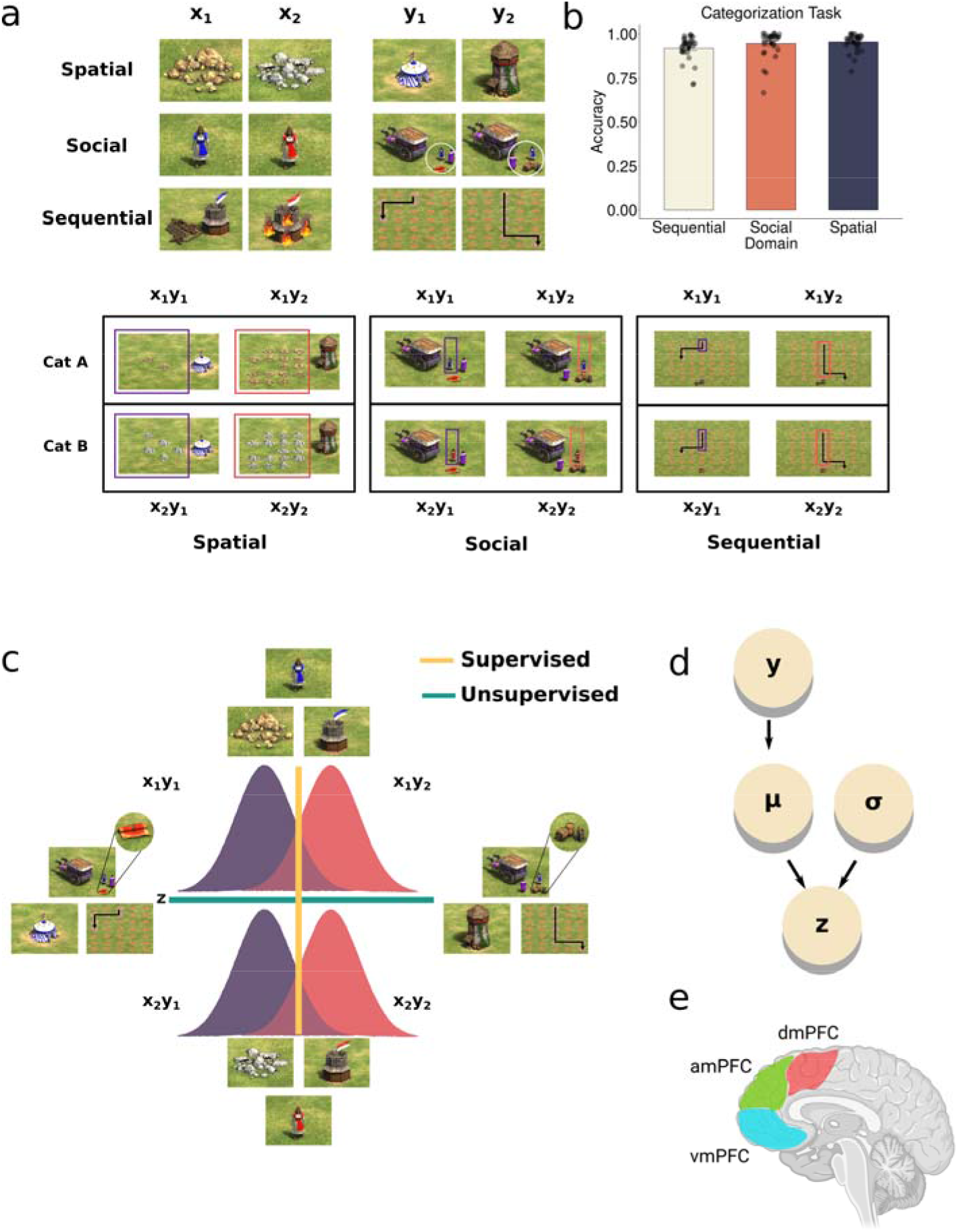
Domain-Specific World Model Learning Task. (a) (Top) During learning, participants categorized exemplars based on an x feature, with feedback, across three different domains: spatial, social and sequential. Stimuli also had a y feature. In order to succeed in a subsequent test phase, participants, in addition to the overt categorization task, had to learn the association between these y features and another continuous feature z (see text). This was drawn from Gaussians parameterized with different means. This resulted in needing to learn four possible ‘states’ of the task based on these features. (Bottom) The z feature was continuous in nature. It represented different properties of the stimuli in each domain but was always drawn from two Gaussian distributions. Smaller z values were typically paired with y_1_ and larger z values with y_2_. (b) Accuracy on the overt categorization task. Dots represent individual participant accuracy and error bars denote standard error of the mean. (c) Full task structure depicting the two orthogonal tasks: An overt (supervised) categorization task based on the x feature and an unsupervised learning of y-z mapping in each domain. (d) Generative structure describing how z features are produced by Gaussians differing in mean parameters, mapped to the y feature. (e) Midline Prefrontal subregions hypothesized to be involved in world modelling.

The common interpretation of amPFC involvement in social cognition is that, like perceptual systems tuned to faces and objects, this region is biased towards processing information in a specific domain of experience: that of social information. However, another possibility is that amPFC is biased towards a particular form of computation that social processing happens to rely heavily on. Specifically, amPFC might be using a general computation of constructing reference frames (i.e., using a coordinate system). Reference frames place features and latent variables onto a common coordinate system to support relational comparisons and goal selection^24,25^. Under this scheme, shifts in people’s mental states correspond to changes in direction within a task-specific basis or frame. Similarly, computing social positions correspond to estimating relations between different points within a given task coordinate frame. This allows comparing social status between novel pairs of people^19^. Indeed, amPFC activity in ^19^ correlated with the magnitude of such comparisons, determined by the direction within the (social) map. Although most commonly linked to social inference, this mechanism may be domain-general, since similar problem structures may occur in other domains. Interestingly, amPFC is involved during tasks requiring maintaining a reference frame^19,26,27^ (or goal) as well as when relational manipulations are required^19,26,27^. This suggests a potential domain-general computational bias for coordinate frames in this region, which social cognition tasks might rely on.

In contrast, ventromedial prefrontal cortex (vmPFC) activity is often observed when encoding spatial associations between multiple variables^28^ as well as in signalling reward valuations^29,30^. These contextual features are assumed to be abstracted into schemas, flexibly deployed during decisions ^14–16^. This domain-specific view suggests that vmPFC encodes value and compacts representations tied to particular sensory cues and spatiotemporal contexts. A domain-general view would be that, rather than coding sensory features or their associated value per se, the vmPFC represents the environmental hidden variables that generate these sensory observations. Concretely, these low-dimensional latent states can be sufficient statistics of a generative model that describes how the sensory stimuli came about. Under this view, classical vmPFC signals (value, schemas, concept abstraction) are consequences of the single operation of locating the current state of the world in the latent space. Thus, inference and selection of states becomes the critical computation here. This provides the necessary mental model for decision-making when key variables are unobserved. Recent results and perspectives also seem to suggest this^7,31,32^, fitting the breadth of vmPFC involvement observed across studies.

Finally, the pervasive involvement of dorsomedial PFC (dmPFC) in action processing suggests a bias towards representing models in the specific domain of action. Here, however, a parsimonious domain-general computational role could be that this region encodes actions across learned latent states, or task policies, whether these are driven by external data or initiated internally^34^. Transitions naturally have a sequential structure and signal specific ways that a given set of states are expected to change under different conditions. This computation is essential for action planning, as actions aim to modify future states of the world in the service of some goal. Predicting upcoming states, and adaptively changing predictions during changes to environment, or changes to one’s goal, are examples of this computation. This domain-general computation could explain puzzling results where dmPFC is implicated in cognitive processes such as mind wandering^35^, surprise/mismatch in one’s predictions (unsigned prediction errors^36^) and foraging tasks^37^, at odds with simple action-oriented perspectives. Mind-wandering involves navigation across internal cognitive states under relaxed constraints. Similarly, planning action sequences involves internally simulating the effects of different strategies or movements across states, under an external constraint. There can be many paths over a set of latent states, each denoting a specific task policy. This requires a monitoring of current policies and their adequacy, possibly via the computation of prediction errors or surprise. Thus, a cross-domain role in computing abstract strategies across states and monitoring them could be the general computational role of dmPFC.

Testing domain-specific vs. domain-general accounts of midline PFC function is difficult for two reasons. First, learning and inference in the domains of state learning, social inference and action are typically investigated in separate studies and are rarely directly compared. Second, when these domains have been studied together, this has often been in the context of naturalistic stimuli, where domains and computations are highly correlated and it is therefore difficult to disentangle their contributions^23,38,39^. In the present study, we address these issues by comparing learning in all three domains and, critically, by orthogonalizing domains and computations.

During fMRI, participants completed three learning tasks, each of which involved learning the mapping between a continuously-varying stimulus feature (*z*) and another stimulus feature that could take one of two values (y_1_/y_2_). Learning happened across spatial, social, and sequential domains: in one task, participants learned spatial associations between two features; in another, they inferred a person’s intention from their interaction with another agent; and in the third, they predicted the action path of a moving object. Each domain thus had distinct surface features, but unbeknownst to participants, all three tasks shared the same latent probability distributions. This design dissociated domain of experience from the underlying latent structure of the information to be learned. If midline PFC activity were domain-specific, effects would differ by task but if it were differentiated by computation, effects would be similar across tasks because the latent structure to be learned is identical in all three cases. We find evidence for the latter and suggest how midline PFC might construct internal models through a set of general-purpose, computations.

## Results

### Virtual World Model Learning Task

We generated 72 virtual-world exemplars using Age of Empires II, spanning three domains associated with distinct cover stories. Each of these exemplars introduced different stimulus features and cover stories (Fig 1a). Before scanning, participants (N=31) completed a pretraining task (see Methods), learning a categorization rule based on one feature (x: x_1_ or x_2_). This feature x was independent of another second orthogonal feature (y: y1 or y2) which was the focus of the main learning task. In the Spatial domain, the features were landmarks and mines (Fig 1a, first row) in a physical environment. Features in Social domain were identity and behaviour (Fig 1a, second row) of two people engaged in a social interaction. Finally, in Sequential domain, the features were trajectories and distances (Fig 1a, third row) of an object moving through space.

Each domain was presented in a separate scanning run. During fMRI, participants categorized the stimuli according to the previously learned x1/x2 rule while receiving trial-by-trial feedback (Fig. 1d). Participants were asked to complete this simple overt task to ensure that they maintained attention and processed each stimulus fully. However, the main focus of our investigation was the covert learning of the mapping of feature y to the continuous property z. Each exemplar had an associated z value which represented a quantity in the exemplar within each domain. These were the number of mines in the environment (Spatial), the speed of a person’s movement (Social) and the distance traversed by the object (Sequential). The values of the z feature were drawn from two Gaussians (Fig 1e). Importantly, each Gaussian was paired with a different y feature. That is, the y_1_ features were associated with smaller values and y_2_ features were associated with larger values (Fig 1c).

Each domain’s continuous z feature and their associated binary y feature formed distinct stimulus sets constrained by the same latent probability distributions. Participants learned these associations in an unsupervised manner^40^. Each exemplar presented could come from one of four categories through a unique combination of x and y: x_1_y_1_, x_1_y_2_, x_2_y_1_ and x_2_y_2_ (Fig 1c). And each one of these could be presented in one of four unique forms based on the z feature. For example, x_1_y_1_ could appear as x_1_y_1_z_1_, x_1_y_1_z_2_, x_1_y_1_z_3_ and x_1_y_1_z_4_ where the values of z were drawn from the smaller Gaussian. These Gaussians described the state of the task at any trial (Fig 1d). A task state here is operationalized as the set of observed features and the unobserved latent distribution thought to have generated them. Overall, the full task involved two orthogonal components (Fig 1c), namely categorizing the exemplars based on x_1_ and x_2_, which formed the supervised part; and separately four mappings of y_1_ and y_2_ features with the z feature, with the knowledge of Gaussians parameters that generated them, which is the unsupervised part.

Thus, although each domain differed in surface characteristics and stimulus properties across the x, y, and z features, all were governed by the same underlying probability distributions defining the latent structure. Following learning, each domain included a test phase to assess whether participants had acquired this latent structure (Fig. 1d). These design features allowed us to adjudicate between competing hypotheses. A domain-specific account predicts that distinct PFC subregions would be sensitive to learning in each domain. In contrast, a computational account predicts similar learning-related effects across domains within the PFC, with differences between subregions determined by the computations they implement rather than stimulus domain. Participants showed near ceiling accuracy in the overt categorization rule in all domains (Spatial M = 95.43, SD= 4.84, Social M = 95.53, SD= 7.64, Sequential M = 92.11, SD = 7.10) (Fig 1b). Importantly, this indicates they were focused on the stimuli and learning task throughout.

### Adjudicating different Representational schemes in midline PFC

We hypothesized that midline PFC regions (vmPFC, amPFC and dmPFC) (Fig 1e) would be representing a model of the virtual world participants are experiencing during learning. We used representational similarity analysis (RSA) to test whether different PFC subregions would represent models specific to each domain (i.e., domain-specific representation).

One way to test this is to see if neural patterns in PFC are tuned to the visible properties of the stimuli that participants encountered during learning. We therefore constructed stimulus-based Representational Dissimilarity Matrices (RDMs) encoding the three feature dimensions (x, y, and z). For each feature, we generated a 24 × 24 RDM representing pairwise dissimilarities between exemplars using 1D Euclidean distance along that feature dimension (Fig 2).

**Fig 2:**
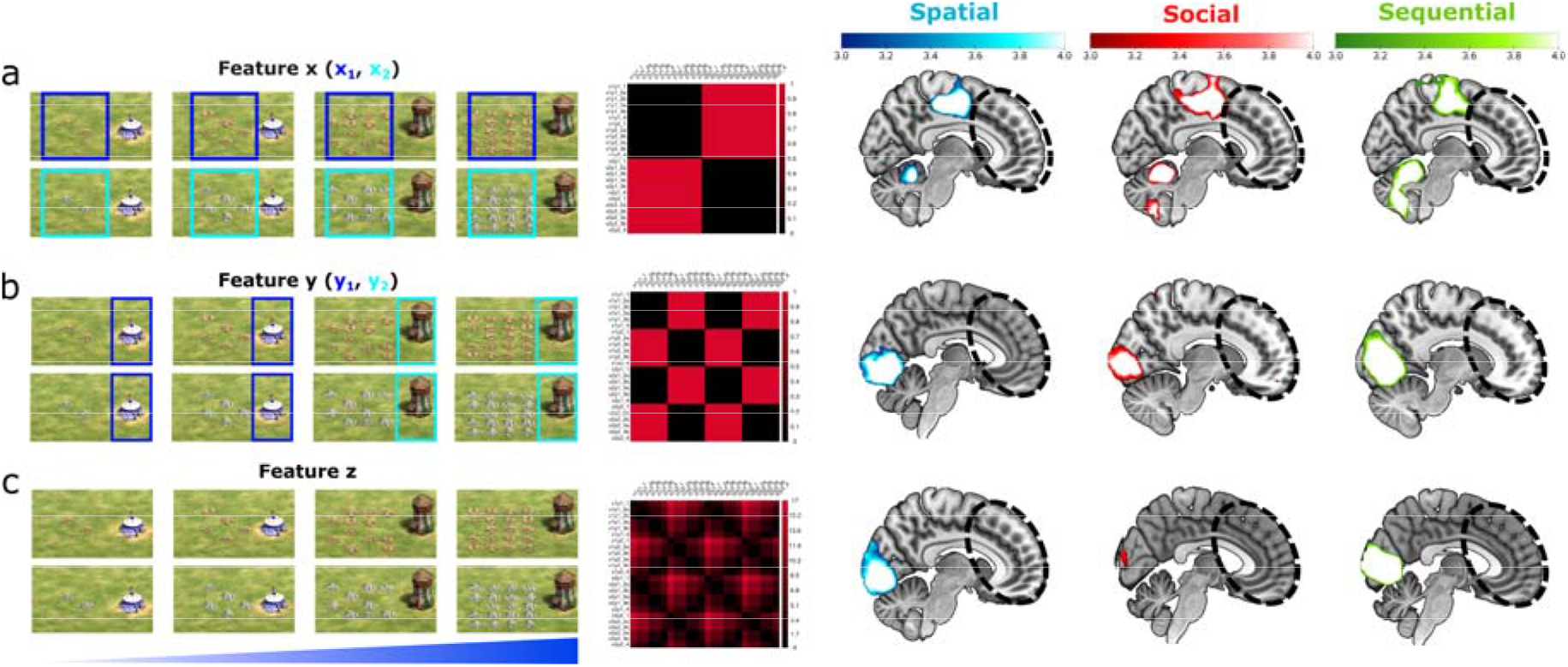
Domain-Specific Feature Representations. Illustration of each kind of representation (left) with stimulus-RDMs (middle) and whole-brain searchlight RSA maps (right) on (from left to right) spatial, social and sequential domains. Spatial domain is shown in a-c for illustration here, but the logic remains the same for social and sequential domains. (a) Stimulus category representation based on x feature identity alone. (b) Feature y RDMs, based on y feature identity. (c) Continuous feature z RDMs generated by calculating differences in the z values between exemplars. No prefrontal patterns were observed to be tuned to these representational schemes across domains during learning (whole brain TFCE corrected p<0.01).

Under a domain-specific account, distinct PFC regions should be sensitive to these stimulus properties in different domains. In contrast, a computational account predicts similar representational effects across domains, given that the underlying computational demands were equated over our three domains.

Thus, we tested whether PFC activation patterns were predicted by (a) the x_1_/x_2_ feature which participants were asked to classify on each trial, (b) the y_1_/y_2_ features or (c) the continuous z feature. All of these are observed properties of the stimuli that are different in each domain. We investigated these using whole-brain searchlight RSA in each domain separately, using multivoxel patterns derived from normalized GLM beta estimates (i.e., t values) from fitting the unsmoothed BOLD time-series (see Methods).

#### Representation 1: x Category

In stimulus category RDM (Fig 2a left, middle), trials with the same x category are coded as similar to each other, independent of their y and z features. This representation is simply the category of the current stimulus learned, formally:

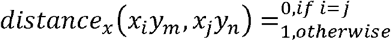

We found category representations in the premotor cortex, primary motor cortex and cerebellum across all three domains (Fig 2a right). These effects presumably reflect coding of the motor responses participants were making during the task. However, this RDM failed to identify regions in the PFC.

#### Representation 2: Binary feature y

We next tested whether PFC encoded the binary y property that was the focus of implicit learning during the task. For this, we constructed an RDM based on the binary feature y (y_1_/y_2_) (Fig 2b, left middle). In this representational scheme, trials with the same y feature are similar to one another while being distinct from trials with the other y feature. This is described as

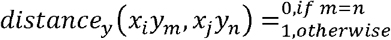

We found significant representational similarity effects in the early visual cortex (V1) (Fig 2b, right) for the y features. Although broadly overlapping regions were implicated, the spatial distribution of effects differed subtly between domains. Spatial domain effects were focused on midline V1. The social domain additionally showed representations in more lateral V1 in the right hemisphere as well as fusiform face area, possibly due to encoding the faces of the agents in this domain. The sequential domain had effects in both midline and lateral V1 in addition to visual motion areas around v5/MT, reflecting sensitivity to the patterns of sequential movement in this domain. Thus, this analysis revealed that the activity in visual regions encoded the features that were relevant for learning in each domain. However, we again found no significant prefrontal representations.

#### Representation 3: Continuous feature z

Finally, we investigated a more continuous representational format, hypothesising that, instead of representing the y feature, the PFC could be representing values of the continuous z feature that predicted it. This RDM defined stimulus dissimilarity in terms of the difference between exemplars on the continuous dimension (z) to obtain a 1D structure (Fig 1c left, middle).

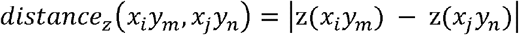

Across all domains, segments of visual cortex showed patterns that reflected the structure in the z dimension (Fig 1c right), again suggesting encoding of visual properties relevant to learning. Despite this, we did not observe significant representational effects in the PFC for this RDM during learning.

To summarise the results so far, we conducted stimulus-based RSA on the expectation that activity patterns in the PFC would reflect the observable features of the stimuli that participants encountered during learning. However, while we found robust effects of stimulus features in visual and motor cortices, we found no effects in midline PFC. In supplementary analyses, we also constructed difficulty-based RDMs to investigate whether midline PFC patterns reflected learning difficulty or effort. These analyses did not reveal PFC effects either (see Supplementary Figures 1-5).

Together, these findings suggest that midline PFC activity patterns did not align with observable stimulus properties or task difficulty during learning. Of course, it is well-known that DMN regions like medial PFC represent *internal* mental models that may be decoupled from current sensory input^11–13,23,41^. Perhaps, rather than representing the overt, perceived characteristics of the stimuli, the PFC represents an internal model of the latent structure of the domain. Since participants are learning throughout the task, this internal model is changing and developing over time, even though the observable stimulus features are not. If this view is correct, it is not surprising that PFC activity patterns are not well-aligned with the observed stimulus features. It also means that to fully understand the role of PFC, we need to predict activation similarities using estimates of participants’ internal models and their trial-wise changes.

Next, we describe the test phase results, investigating how participants learned the tasks across domains. We used these behavioural results to validate a computational model of learning in our tasks. We then used this model to generate estimates of participants’ latent task representations at different points in learning. Finally, we were able to use these estimates as predictors of neural activity.

### Testing World Model Learning Across Domains

We next used the behavioural responses collected during the test phase to assess different models of how participants learned. During the test phase, participants sequentially observed two exemplars in which the z feature took on novel values not encountered during learning. The y feature was hidden and had to be inferred based on the value of z features (Fig 3a). Participants were informed that both exemplars within a trial had the same (unobserved) y feature. Optimal inference therefore required them to have learned the mapping from z to y (i.e., which Gaussian could have produced the z feature) (Fig 1d). Test exemplars were drawn from different parts of the distribution and in different orders, resulting in seven test pair conditions (Fig 3b). Some exemplar pairs were taken from the extreme ends of the two Gaussian distributions (labelled E1, E2), which are strongly associated with one of the two y features. Other exemplars were taken from the central region in which the two Gaussians substantially overlapped. These exemplars were not strongly diagnostic of the y feature, so they are labelled A1, A2 (for ambiguous). There were also other combinations based on selecting one from each of these types (e.g., E2A1) (Fig 3b). We are thus able to vary the difficulty of inference in the test phase. On EE trials, it was easy to infer the correct y feature. EA trials made it relatively more difficult since the A exemplar might favour the incorrect y feature (A2>A1). Pairs were presented in different orders (Fig 3b).

**Fig 3:**
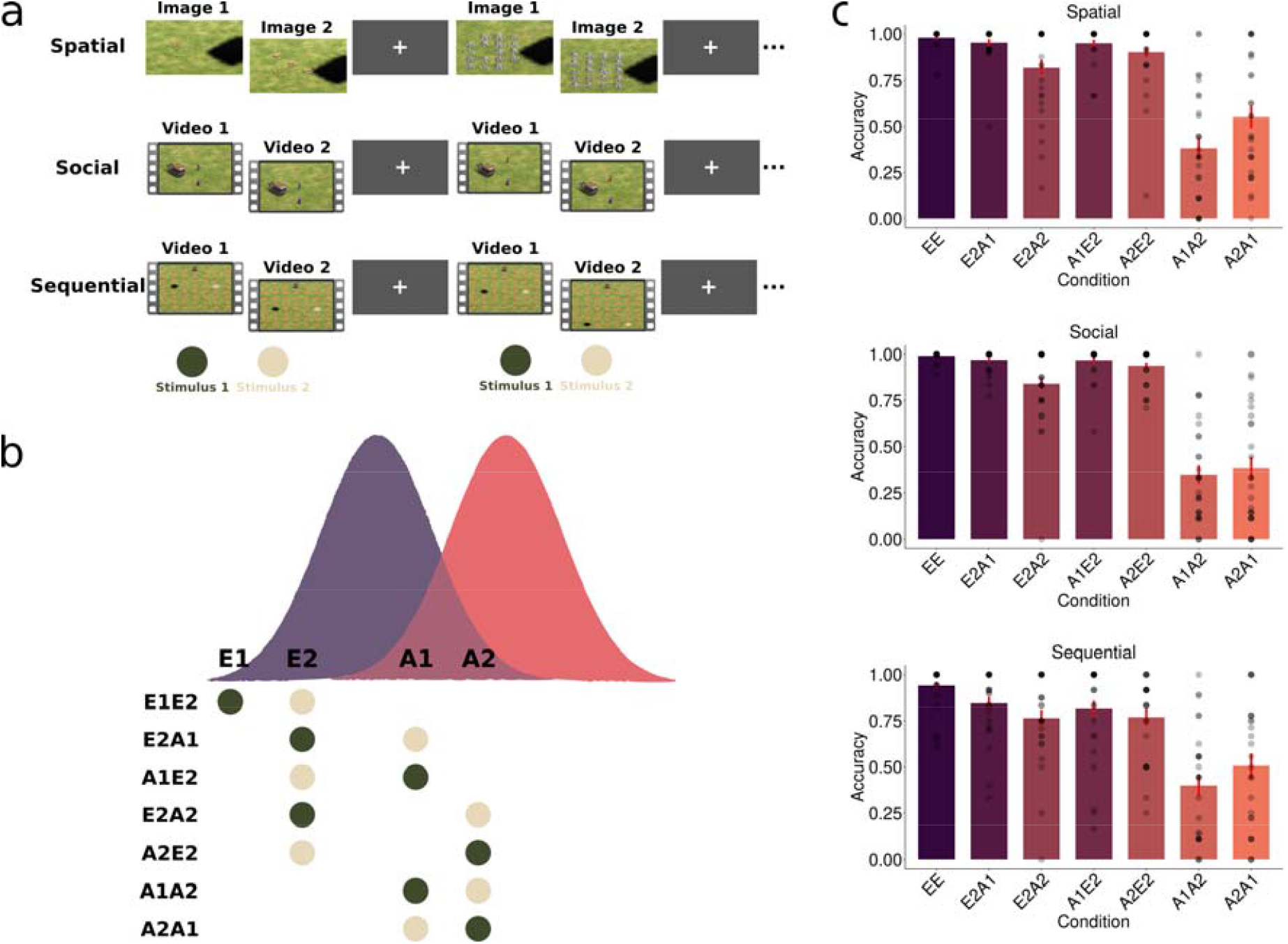
Test Phase Task Structure and Behaviour. (a) During test phase participants saw a sequence of two test exemplars (e.g., E1E2), drawn from the same x category and y feature combination, but with novel z feature values and for which the y feature is now occluded. Participants had to guess whether the occluded feature was y_1_ or y_2_ (e.g., tent or tower in spatial domain). (b) Test exemplars were drawn from different parts of the distribution and in different orders, resulting in seven test pairs. These included an easy scenario with a pair of exemplars from the extremes of the gaussians (E1E2), fully ambiguous pairs (A1A2) and combinations thereof. (c) Test phase performance over each domain split by condition. A1A2 and A2A1 trials are ambiguous and do not have a strictly correct choice. For visualisation these trials have been coded with y_1_ as the correct option. Dots represent individual participants and error bars denote standard error of the mean.

We compared the test accuracy across the 7 test conditions. AA scenarios did not have a correct answer so to visualise responses on these trials, we arbitrarily chose to code y1 responses as correct and y2 as incorrect. An unbiased learner would score 50% on these trials; deviations from 50% represent a consistent bias towards one or the other y feature. Overall, participants had high accuracy in the EE scenarios across domains (Spatial M=92.4, SE =4.07, Social M = 99.2, SE = 0.4, Sequential M = 94.2, SE =2.3) (Fig 3c). The AA scenarios often differed from 50%. Moreover, they varied across domains, showing some order effects (A1A2 vs A2A1 Spatial p = 0.003, d =0.59, Social p = 0.45, d =0.14, Sequential p = 0.13, d =0.30) and also exhibited high variability across participants (Fig 3c). These results suggest participants were consistently biased towards one distribution in fully ambiguous scenarios. However, the size and direction of the bias varied widely among them, suggesting within-subject biases rather than a simple group-level preference towards one y feature.

In the EA pair scenarios, two domains showed a notable reduction in accuracy when the A (ambiguous) exemplar seen had a higher likelihood under opposing model (E2A1 vs E2A2 Spatial p = 0.006, d =0.56, Social p = 0.0009, d =0.68, Sequential p = 0.17, d =0.27) (Fig 3c). Participants were also less accurate on trials where they saw the high likelihood exemplar (E2) first followed by the ambiguous ones (A1 or A2), compared to the reverse but this trend was not consistent (A2E2 vs E2A2 Spatial p = 0.29, d =0.20, Social p = 0.03, d =0.42, Sequential p = 0.93, d =0.01).

These results suggest a sequential order effect as well as bias towards one of the distributions by different participants, irrespective of the domain. We next illustrate how an internal model could be constructed and examined whether its internal representations can account for the observed behavioural patterns.

### General Computations for World Model Learning

Broadly speaking, there are distinct computations that may be involved during formation of internal models from observed sequences of data. The goal of the participants was to infer the parameters of the underlying Gaussians from only the z features, in a trial-by-trial manner (Fig 4a). Later during the test phase, they would select between competing Gaussians from only two novel z features, drawn from various zones (Fig 3b). An optimal way to do this in our learning task is to encode the Gaussians, which cannot be observed and thus needed to be inferred probabilistically. This reduces to a problem of Bayesian inference where participants might use a prior belief about the latent parameters of the Gaussian, and updates them sequentially as they observe more trials (Fig 4a, top). This results in a posterior belief over the parameters of the Gaussian, which becomes their prior for the next trial, and so on. Exact Bayesian inference is thought to be intractable^42,43^ for the brain to implement due to resource constraints; therefore it is often assumed that people use approximate inference^44,45^. One candidate form of such approximation is Monte Carlo sampling. Sampling-based algorithms have been shown to explain human higher-order cognition better than optimal, normative flavours of Bayes. There are many varieties of sampling, but here we use Markov-Chain Monte-Carlo (MCMC) as a model of how the brain might explore the parameter space during learning phase (Fig 4a, bottom). We make no assumption that this is the way the brain necessarily implements probabilistic inference, but only that it is a form of approximation that is well-suited to the task goal, with evidence from a number of behavioural^45,46^ and theoretical^47,48^ results, in similar settings of latent variable inference.

**Fig 4:**
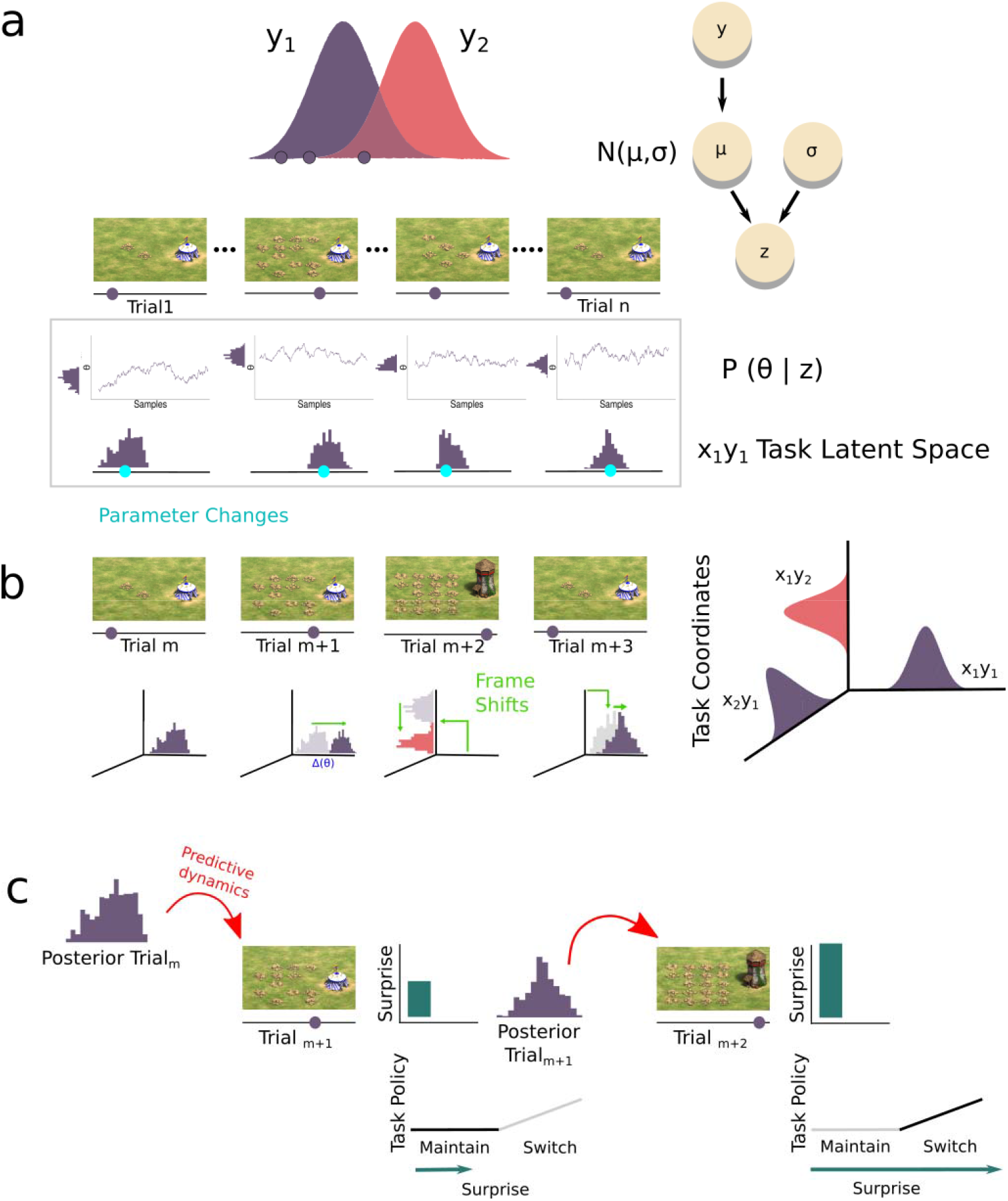
Specialized Computations for World Modelling Schematic. (a) (Top) Learning exemplars were drawn i.i.d from their respective Gaussians. Each exemplar has a particular z feature value (here the spatial domain is shown). (Middle) Each trial, a Markov chain is initialized which searches (locally) and updates (incrementally) the mean and standard deviation parameters, which together represent the Gaussian density (only showing the mean parameter chains here). Y axis represents the mean value and X axis denote the number of samples within a trial (Bottom) They can be visualized on a latent space jointly representing the posterior beliefs about µ at any given trial (cyan, denoting the posterior expectation). This initially starts with wide, uncertain beliefs about parameters and as learning progresses approximates to true values with more precise beliefs. Importantly, these values exist in the (parameter) latent space and not in the stimulus space. This can be intuited by looking at the posterior in trial 1 and trial n, both of which have the same stimulus z feature (2 mines) but have different latent representations. Conversely, trials shortly before trial n have similar latent space locations despite different stimulus values. This suggests the learner has encoded the posterior location that describes the Gaussian that has generated the data, or the state of the task. (b) There are four such latent spaces, representing each category/state, which are orthogonal to each other (for visualization only three are shown with x_2_y_2_ assumed to be in another axis). Multiple task states could be organized along orthogonal basis frames for a single global task coordinate system with directional changes (light green). These can change direction depending on the updates in the latent space as well as shifts onto other axes when states change in learning (c) (Left) Current posterior has a predictive density (see Main text) that assigns future z values (trial m+1) probabilities given the posterior (trial m), which results in surprise values (dark green) for each exemplar through its forward predictive dynamics (red). A task policy unique to a state which was maintained after observing trial m+1, (low surprise) is then changed on trial m+2 (high surprise), incurring a cost in behavioural responses.

This computation results in a posterior over Gaussian parameters, inferred from the z features and which is conditioned on the observed x and y feature, resulting in a task state (Fig 4a). We operationalize this as task state because it encapsulates the full information of the learning task at that given trial, namely the x feature (required for the category response), y feature (required for learning) and the parameters comprising the Gaussian, from which the z feature was likely generated. Knowing the parameters of the state is critical to infer the correct y feature during the test phase. This computation may enable an internal model to approximate the latent task state by exploring the parameter space through a sampling process, rather than maintaining full probability distributions. Thus, estimating the task state is the first computation of an internal model in our design.

However, in our task there are four such states that participants encounter during learning, namely x_1_y_1_z, x_1_y_2_z, x_2_y_1_z and x_2_y_2_z. This is because x_1_ and x_2_ are category-coded orthogonally, even though the same Gaussians are used in each. In addition, y_1_ and y_2_ are also orthogonal in the test phase to be adjudicated. This suggests a mechanism to represent these four states separably could be used. If so, this aspect of the internal model could help shift the participants between the four states adaptively between them according to the trial information of x and y. For example, consider if one state (e.g., x_2_y_2_z) was encountered much later than other states in the trial sequence. Its parameters need to be inferred without interference from similar-looking states (e.g., x_1_y_2_z, which has the same Gaussian density but is a fundamentally distinct state) which at that point might have a different posterior. Representing these relevant task states as separate, and switching between them is the second computation. A parsimonious way to achieve this is to construct a task coordinate system (Fig 4b). Much like how semantic information is arranged on a high-dimensional coordinate system with different features along separate axes, here the abstract states could be arranged along independent axes. In parallel to the probabilistic computation for state estimation, this geometric computation of forming a reference frame that organizes and tracks each state as it updates, and switches between these states adaptively, is crucial in our design.

Beyond tracking latent state parameters, world model learning also requires evaluating whether the current strategy (or task policy) for acting within a state remains viable. A policy is a specific way of taking actions within a state. We propose that participants maintain a forward predictive model by computing the likelihood of each incoming observation under their current posterior belief (Fig. 4c, left). Although the randomised trial structure renders explicit prediction non-instrumental, this surprise signal provides a behaviourally consequential index of how well a strategy accounts for unfolding observations. Policies must be maintained with sufficient stability to support consistent behaviour, yet remain sensitive to evidence that a strategy is no longer adequate. Surprise^49–52^ provides a natural signal for this evaluation: high surprise indicates the current policy is being violated by incoming data while low surprise indicates the trajectory is proceeding as expected and the policy can be maintained.

This third computation, forward predictive evaluation of task-state dynamics for policy tracking, is dissociable from state inference and coordinate organisation. Together, these three computations constitute a division of labour where the first one maintains the evolving task state parameters, the second organises states into a task coordinate frame, while the third instantiates and monitors whether the current policy over those states remains adequate.

In order to test how these computations are instantiated in PFC, we needed to obtain estimates of the latent state parameters on each trial. Next, we evaluate candidate learning models that could provide these estimates.

### Adjudicating different Learning schemes for World models

We considered several different ways that participants might encode information required to perform successfully in the test phase, ranging from encoding a simple decision boundary^53,54^ to memorizing exemplars (prototype model)^55,56^, to forming a complete distributional representation of P(z|y) (Fig 5a). Alternatively, some participants might not have encoded the required information at all during the learning phase, leading to responding randomly in test phase. Our first three models of learning simulated these situations: a Random Responder model, a Boundary Learner and Prototype learner^57^ which encodes the conditional mean of the z values associated with the y feature, but not the full distribution. For the Prototype learning, test pairs can be compared to the mean of each p(z|y_1_) or p(z|y_2_) to make a choice (Fig 4a). Then there is inference to the generative distribution p(z|y) for which one should in this case ideally infer a posterior on the sufficient statistics of the two y-conditional Gaussians e.g. [μ_z|y_1_, μ_z|y_2_, σ^2^_z|y_1_, σ^2^_z|y_2_]. (See Methods for detailed descriptions of these models).

**Fig 5:**
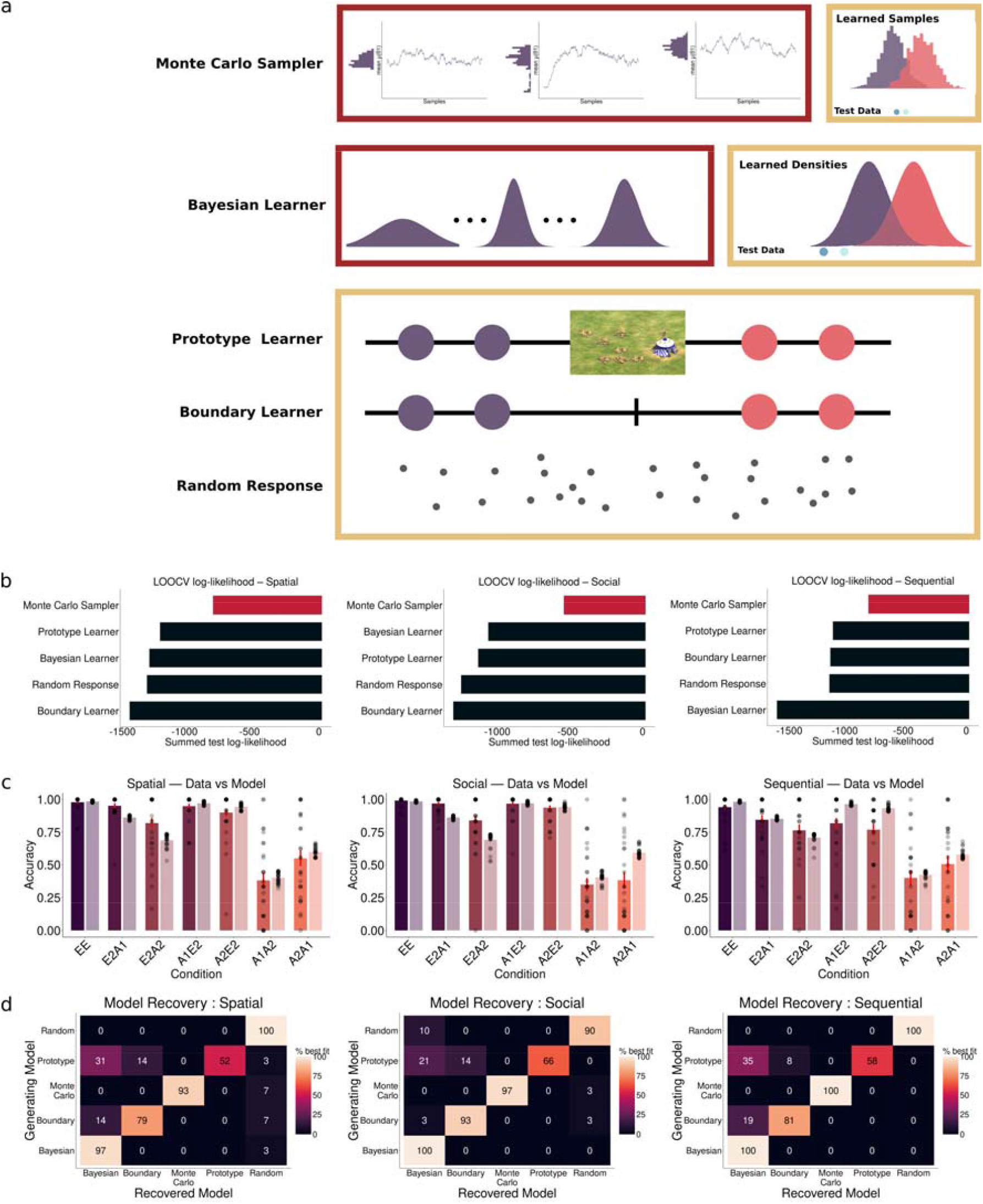
Computational Modelling of various Learning Strategies. (a) Various Learning strategies participants could deploy to learn the latent structure (see main text for description). (b) Leave-one-condition-out cross validation across the models for assessing model fits. A Monte Carlo sampler had the highest loglikelihood for explaining participant behaviour across all domains. (c) Fits by the model (fitted Monte Carlo sampler) (light) and participant behaviour (dark) across all seven conditions. A1A2 and A2A1 trials are ambiguous so their response is coded as proportion of y1 response, as they do not have a strictly correct choice. Dots represent individual participants and error bars denote standard error of the mean. (d) Confusion matrices from a model recovery analysis using simulated data from fitted models showing most learning strategies were well distinguishable from one another.

Our final two models took a Bayesian approach to the challenge of learning. In the normative Bayesian learner, for each of the four category-context pairs (x_1_y_1_z, x_1_y_2_z, x_2_y_1_z, x_2_y_2_z), we assumed participants modelled the latent structure as a Gaussian with an unknown mean μ and variance σ^2^. This is because the Gaussians determine the mapping of z values to y feature. A Normal-Gamma prior over the parameters was used (with weak hyperpriors), allowing closed-form, single-trial updates during learning^58^ (see Methods). Thus, after every learning trial the model had a posterior over parameters. At test, one can then calculate the posterior-predictive likelihood of each test pair by integrating over these parameter posteriors. A soft-max with a temperature parameter then converted the posterior into probabilities of responding y_1_ or y_2_. In this benchmark model, we assume a) participants had inferred posteriors on y-conditional means and precisions, by performing b) closed-form analytical updates but c) deviated from hard maximization over this posterior due to decision noise.

The final model is the Monte Carlo sampler which performs approximate Bayesian inference. We formalized an online MCMC sampler to this end. During learning, after each categorisation trial, the sampler treats the observed z values as draws from an unknown Gaussian distribution. It estimates a posterior distribution over the parameters of this Gaussian by generating a chain of samples from p(μ_z_, σ^2^_z_ |z_1:t_) based on the trial history of observed z values; we here simplify notation μ_z_ and σ^2^_z_ to μ and σ^2^ henceforth (see Methods for full description). The sampler, instead of storing past data (z_1:t-1_), was based on sufficient summary statistics: how many observations have been seen, the sum of all values so far, and the sum of their squares. Note that although these quantities are sufficient, and the relevant posteriors on μ and σ^2^, and predictive likelihoods for new z samples are analytically calculable in this specific instance, this is not generally the case^45^. Thus, we model the parameter inference (μ and *σ*^2^) is still a search process i.e., sampling from the posterior. The sampler starts with weak, uninformative priors and distributions on all parameters are approximated via chains of autocorrelated samples. Rather than closed-form computing of the posterior as in normative Bayes, here it is approximated via Metropolis-Hastings. After every learning trial, the sampler recursively estimates the posterior by using the likelihood with updated sufficient statistics. we later use the mean of each of these interim posterior samples to construct RDMs that capture the internal models of each participant, across trials.

Humans often behave as if they are basing decisions on a relatively small number of samples^44^, often just a single sample^59^ or computing simple summary statistics^60^ of these samples, to balance computational costs against the benefits of additional precision. We formalized both these ideas within our sampler, hypothesizing that approximation by using only a limited set of samples^46^. through a likelihood-free inference scheme^61^, also known as Approximate Bayesian Computation (ABC). Rather than performing model selection by evaluating the full closed-form likelihoods of the test exemplars under each state as p(z|y_1_) and p(z|y_2_), it instead computes the distance between test z-features directly to the internal samples generated from the learned posterior (see Methods). Samples are accepted when their distance from the observed data falls below a tolerance threshold. The resulting relative acceptance rates of the samples from p(z|y_1_) and p(z|y_2_) are used as a proxy for the categories’ relative likelihoods (Fig 7a). The tolerance, the model’s only free parameter, therefore determines how permissive this matching process is. Higher values allow more simulated samples to be accepted, whereas lower values enforce stricter matching. Moreover, we assume the posterior samples recruited for evaluating the first test exemplar are then reused (after normalizing) as prior samples for the second test exemplar, naturally producing sequential effects (Fig 7a). The sampler thus predicts participant-specific biases that depend on tolerance value and generate distinct order effects even within conditions with same z-features, such as A1A2 versus A2A1. This is because the second-stage inference depends on how many samples survive the first-stage filter.

**Fig 6:**
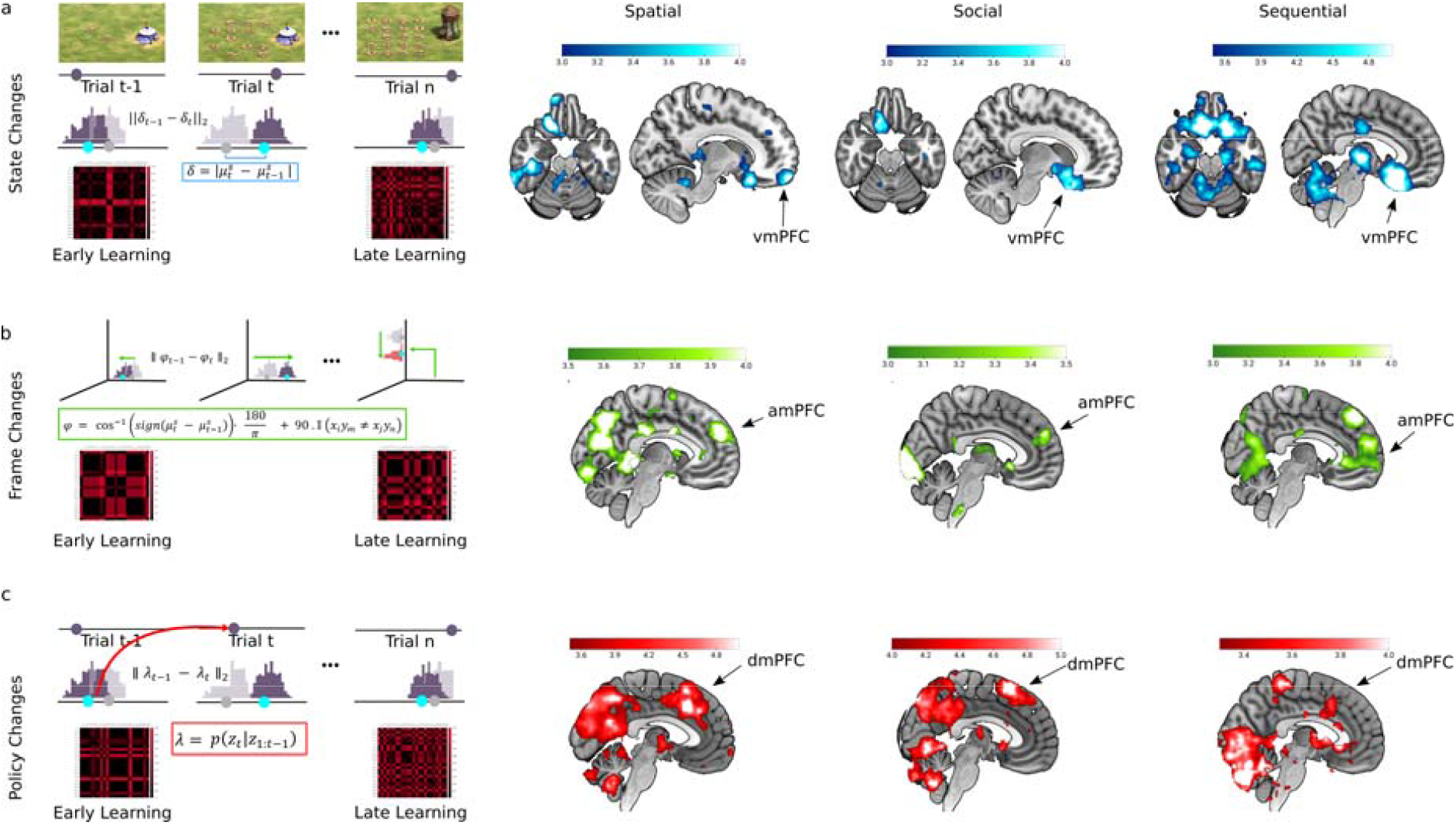
Specialized Computations for World Modelling within the Prefrontal Cortex. Illustration of each kind of computational variable (left) used for a process representation (middle) with model-derived RDMs from a representative participant (below) and whole-brain searchlight RSA maps (middle) for (from left to right) spatial, social and sequential domains (TFCE corrected p<0.01). (a) State changes denote a variable delta δ that registers how much a posterior location in the latent space has changed, and dissimilarity between trials based on this value is used to construct 24x24 RDMs for RSA. RDMs depict early learning vs late learning with less difference between trials during late learning (values are smaller in late learning RDMs). This variable explained the vmPFC neural patterns across domains. (b) Frame changes, [l], denote whether a change in reference frame occurred due to local directional shifts between two posterior locations in subsequent trials within a state as well as global shifts between states (see main text). These RDMs explained amPFC neural pattern changes across domains. (c) Transition changes, λ, denote dynamics of the observed z feature, based on predictions under the posterior predictive and suggested how task transitions can occur. Similarity in transition changes across trials could explain the dmPFC voxel patterns across domains.

**Fig 7:**
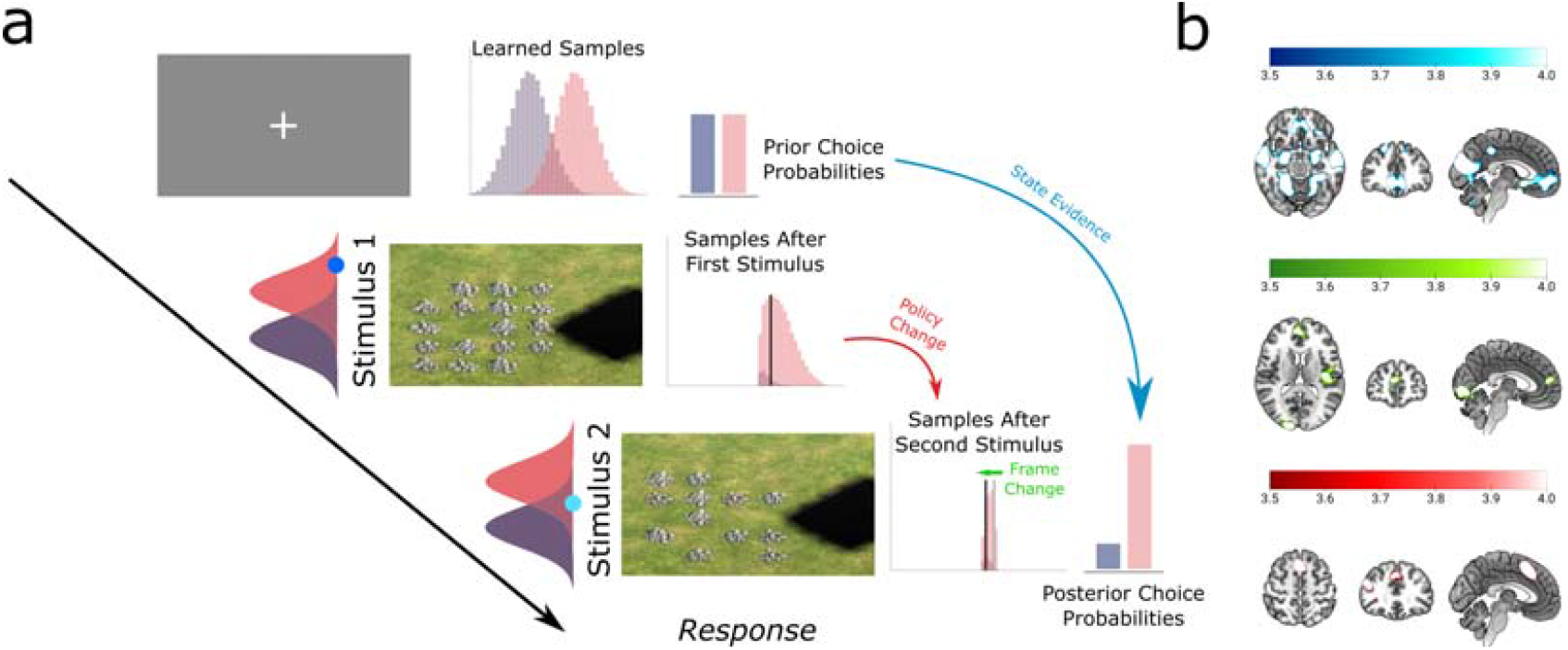
Specialized Computations during Inference within the Prefrontal Cortex. Whole-brain parametric analysis using regressors derived from Monte Carlo sampler during test phase. (a) On each trial, starting with equiprobable choice, participants simulate from their learned samples, comparing internally generated simulations (see main Text, Methods) to test data. They do this sequentially through two exemplar stimuli presented within a trial, reusing the posterior from one stage as the prior for the next. This process involves the same three computations investigated during learning. (b) (Top) State evidence for one state over the other, resulting in their responded posterior probabilities within test trials parametrically modulating vmPFC activity. (Middle) Frame changes within trials modulating amPFC activity. (Below) Policy changes within test trials with more surprising transitions modulating dmPFC BOLD activity.

We compared the above five models to each participant’s test data in each domain through leave-one-condition-out-cross-validation (LOOCV) by training on all conditions minus one, maximizing the likelihood of model parameters generating the data, and then testing with these parameters on the held-out condition. Out-of-sample test predictions were then used to compute total log likelihood for each model as a measure of model fit. Prototype and Decision boundary models only required the test phase data to fit as they assume that participants encoded the true mean values in learning. The Bayesian learner and Monte Carlo sampler, in contrast were exposed to the learning trials. Thus, for each participant, these models entered the test phase with a posterior based on the summary statistics of all learning trials.

In each domain, we found that the Monte Carlo sampler could explain the behavioural data better than other learning strategies (as indicated by smallest negative log-likelihood values in Fig 5b). In addition, it uniquely reproduced participants’ behavioural patterns including biased responses and sequential effects (Fig 5c).

Interestingly, we found that different participants used different tolerance across domains. Specifically, the sequential domain had higher tolerance (M = 4.8) compared to spatial (M = 4, p = 0.03, d = 0.58) and social domains (M = 3.8, p = 0.007, d = 0.75) (Supplementary Fig 6). This may suggest a task policy where an increased tolerance is implemented to control inference over different domains.

Moreover, both the Monte Carlo sampler and normative Bayesian Learner could be differentiated using model recovery analyses, suggesting they both produce distinct behavioural profiles (Fig 5d).

The modelling results above suggest two key results. First, a Monte Carlo sampling-based process, which used a comparison of internally simulated samples to externally observed data, could capture and reproduce human-generated behaviour more accurately than normative Bayesian, and non-probabilistic heuristic models. Particularly, the reliance of output of an internal model and the reuse of these samples underpinned the sequential effects observed in human behaviour. Second, this was consistent across domains suggesting a common process for learning latent structure by approximating Bayesian inference. Next, we investigate the neural correlates of these different internal model computations involved within this process.

### Testing Neural Representations of World Model Learning

In the first section of this paper, we tested whether PFC regions encoded the observable features of stimuli while learning in three distinct domains. No evidence of such coding was found, leading us to hypothesise the PFC may instead be representing internal models of the latent structure underlying each learning task. The Monte Carlo sampler we have just described provides us with estimates of the status of these internal models during learning, on a trial-by-trial basis, for each individual participant. This provides us with an empirical basis for seeking neural correlates of internal model formation. In the remainder of the paper, we outline three different ways in which neural processing might act upon internal models in our task and then test for evidence of such processing in PFC. There are at least two ways in which processing in PFC might be organised. On a domain-specific view, specialisation is determined by the domain the stimuli belong to (spatial, social or sequential), with no differentiation based on the type of computational processes. Alternatively, a domain-general view would predict specialisation for different computational processes, no matter which domain learning is occurring in. Our results support a domain-general organisation.

### Domain-General Latent Task State Representations in vmPFC

As described before, the most fundamental computation required for modelling the world is to accurately infer and update the states that generate the observed data (here z feature, conditional on the x/y category). The task state at any trial is a category, with corresponding Gaussian that describes how the z feature in the (x,y,z) triplet is generated (Fig 4a, right). Learning the posterior parameters of this Gaussian conditioned on x,y,z results in knowledge of the task state parameters as p(μ|z,x,y). There is considerable evidence that the brain organizes abstract task and conceptual spaces in geometric ways akin to physical space, known as cognitive maps^14,62^, allowing for generalizations^19^ while also being structurally compact^63^.

Our Monte Carlo sampler assumes that participants used a stochastic search process to approximate distributions on the (task) state parameters with a set of samples from P(μ_z, σ^2^_z |y). For a given latent state (x_i_y_j_z), learner maintains a posterior over the Gaussian mean parameter *μ*. The mean parameter changes from trial-to-trial as it gets updated, converging closer to the true data distribution (Fig 4a, bottom). Importantly, this occurs in a latent space of parameters independent of the observation space (Fig 4a, Fig 6a left). As an example, for the state (x_1_y_1_z), early in learning, this posterior can shift substantially from one trial to the next. For instance, suppose that after observing a feature value z-feature = 2, the posterior mean was estimated as μ = 4.5. On the next encounter with the same state, observing z-feature = 12 updates the posterior mean to μ = 9 (Fig 6a, left, Trial t). The magnitude of belief update is therefore Δµ= 4.5. Later in learning, updates become smaller as the internal model stabilizes. For instance, the posterior mean may change only slightly across successive encounters with the same state (e.g., going from μ = 6.9 and μ = 7.0), yielding Δµ= 0.1.

These trial-by-trial changes in μ define a one-dimensional trajectory through an internal belief space. Each trial thus corresponds to a location on the μ-axis, and learning corresponds to movement along this axis. We therefore tested whether PFC activity patterns track the magnitude of this movement, Δμ, as a proxy for the changes to participant’s internal model.

We used RSA to investigate whether this computational process is reflected in PFC activity patterns. For this we derived RDMs from each participant’s learning model. We obtained the model parameters for each participant’s Monte Carlo sampler (fitted to their test data). Then we submitted to the model the sequence of trials observed by that participant during learning. This generated trial-by-trial posterior parameter values (μ) for each participant for each trial they experienced in the scanner.

Next, for each trial we calculated how much the posterior mean changed compared with the previous trial, Δµ, for each state (using s here as a subscript for ease of use over x_i_y_i_z):

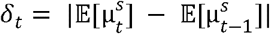

Here, the 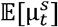 denotes the expectation over the posterior mean samples at trial *t* under a unique state, *s*.

We constructed RDMs based on these values (Fig 6a, left column). Here trials are similar if they change posterior locations to a similar degree relative to the preceding learning trial.

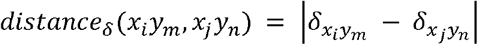

Example RDMs are shown in Fig 6a left. In early learning blocks the representational distance between trials are relatively high (Fig 6a, left column). This is because the participants have only rough estimates of the distributions, and parameters progressively converge within each state during later learning blocks.

We used three whole-brain searchlight analyses (one per domain) to test for regions whose activation patterns were predicted by similarity in the posterior change. Within PFC, ventromedial PFC pattern representations showed high similarity with how each participant’s posterior beliefs changed (Fig 6a, right). Notably, this effect was consistent across domains, demonstrating a domain-general encoding of posterior beliefs across the latent parameter space in the vmPFC. The vmPFC has been observed to encode such task maps in low-dimensional forms^7,15,16^, but the process by which this occurs remained unknown. Our data suggest that it infers latent states through a domain-general probabilistic inference, which argues against domain-specific encoding. The analysis also identified distinct parts of the anterior temporal lobe which varied across each domain (Supplementary Fig 7-9).

This shows two important results. One, representations in the vmPFC might not be encoding domain-specific features of the environments but rather a domain-general acquisition of the states and state parameters that could have generated these observed features. Two, such a process representation is unique to the learning trajectories of each participant and implicitly compresses the distribution to the task state.

### Domain-General Coordinate System for State Navigation in amPFC

The above results showed that changes to task state location in the latent space described the vmPFC patterns, corroborating a number of studies^14,15,19,28,64^ suggesting geometric coding of abstract spaces in this region. This geometric interpretation of learning task states implies that updating one’s current beliefs of the state involves movement within this latent space. In our task, there are four distinct states to keep track of (x_1_y_1_z, x_1_y_2_z, x_2_y_1_z, x_2_y_2_z). Even though the same Gaussian parameters are used on x_1_ and x_2_ trials, the nature of the overt categorization task makes it necessary to represent these orthogonally. Moreover, y_1_ and y_2_ are also learned separately as they would be tested later on. Our experimental task structure therefore has four orthogonal states and thus four distinct latent spaces (Fig 4b). Participants learned each state without necessarily knowing they come from the same Gaussian distributions. They also did this in a piecemeal way observing one z feature at a time. As such, keeping track of prior locations in each latent space becomes important, both to encode prior task state locations and to minimize interference between these. These call for a second computation of keeping a persistent and stable task reference frame.

As described before, participants might be updating their parameter μ in latent space for a given state, and there are four such states orthogonal to one another (Fig 4b, left and also Fig 6b, left). Take the previous example where at a given trial for the state (x_1_y_1_z), the posterior mean was estimated to be μ = 4.5, is now updated to be μ = 9 in the next trial (Fig 6b, left, Trial t, and Fig 4b, left, Trial m+1). The resulting change in mean is now a signed change of +4.5 and directed to the right. The direction of this update here is 0-degree (as sign change is positive) along the latent space. Consider that the next trial has a different state (x_1_y_2_z) which was also updated from its previous instance some trials before (Fig 6b, left, Trial n and Fig 4b, left, Trial m+2). Firstly, the orthogonal representation would mean that a shift to another orthogonal axes is performed, resulting in a 90-degree change. Secondly, if its previous μ is now, for example, updated from 15.5 to 12.5 the difference is −3 now (sign change is negative). This results in a further 180-degree shift along that axis in the opposite direction as for that state (Fig 6b, left, Trial n). Maintaining such a reference frame helps adaptively switch between states and carry on the direction of updates when state parameters are updated.

Generally speaking, taking the angular change of sign of Δµ (i.e., change in posterior mean) gives the direction of the update in this one-dimensional space: 0° versus 180°. This captures a local reference frame within a latent state. In addition, we assume that the four latent states share a global coordinate system, with each state aligned orthogonally to the other three (Fig 4b). Under this account, switching between latent states produces an additional 90° rotation. Taking these two components together, we define the total angular change as the sum of the local update angle and any global shift between latent states:

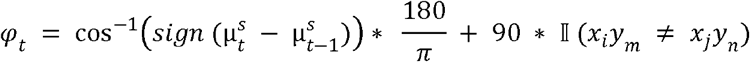

The RDM we constructed for this property codes trials are similar if they encode similar directional shifts, with a minimum 0 degree (exactly same frame changes) to a maximum 270 degrees (where one trial had both local and global shifts, compared to the other) (Fig 6b, left).

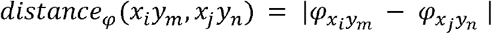

We used three whole-brain searchlight analyses (one per domain) to test for regions whose activation patterns were predicted by similarity in direction change. We found that the anteromedial prefrontal neural representations were explained by changes to the global coordinate shifts in all three domains (Fig 6b, right).

The consistent involvement of amPFC in coding direction shifts across all three domains suggests that this region may encode directional computations for representing reference frame estimations in latent spaces. One important consequence could be that instead of encoding states of the task, this region might be providing task-appropriate reference frames to be used by other regions, notably the vmPFC.

### Domain-General Task Policy Dynamics in dmPFC

Learning a world model also entails a prediction of future environmental states or a representation of future dynamics given past history. In our task, we assume after every trial, given the features and the state parameters learned so far, participants would be making predictions on upcoming trials in a probabilistic manner or keeping a running history of recent experience (Fig 5c). Although in our task such predictions are not helpful (since trial orders are fully randomized), this would result in quantifying how surprising is a new observation given previous estimate of the posterior beliefs. In the general case, this helps adaptively changing task policies (i.e., what actions to be taken at each state) in the face of error.

As an example, suppose after a given trial for state (x_1_y_1_z), the posterior mean is estimated at μ = 9 (σ = 4). The next trial, upon observing z = 12 (Fig 4c, left, Trial_m+1_), the model registers a surprise proportional to the inverse likelihood of that observation. In isolation, this is a prediction error. But in the context of a maintained task policy, it is also a signal about whether that strategy should continue. If in the next trial if the observed z = 19 and y feature is different, it indicates a switch in latent state (x_1_y_2_z) thereby suggesting a change in policy. A high surprise thus would indicate the policy needs revision, in learning and test phase. Crucially, this signal has direct behavioural consequences. Across domains, higher surprise during learning slowed response times within a trial as observed from a linear mixed-effects model (β = 0.002, p < 0.001) (see Methods). The same pattern held at test, where surprise continued to modulate response times (β = 0.03, p < 0.001). Importantly, this is in the absence of further or minimal learning, indicating that the policy of choosing the states was being instantiated and actively evaluated against novel observations reflected in a costly increase in reaction times. We investigated then, whether the surprise signals that lead to the task policies were represented in the PFC during learning phase.

The formal way of characterizing probability of newly observed z features, given the observed history so far is given by

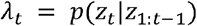

This is the posterior predictive at each trial. This can be expanded as

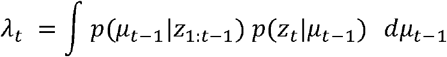

Where the first term within the integral denotes the posterior means (*μ*) learned until the previous trial, given the z features observed as far as then. The second term denotes the likelihood of new z features under the previous posterior mean, integrated over all possible values. In other words, this is a predictive density of new z features given what participants learned so far. A high value suggests that the observed z feature in current trial had the highest predicted probability under the previous posterior belief. Low values, in contrast denotes highly surprising values or low probability, again under the previous posterior mean of the state. Surprise, here is the inverse of this where low values quantify higher unexpectedness.

Trials that had similar surprise values would be more similar to one another (Fig 6c, left). RDMs were constructed based on this (Fig 6c, left); these were unique to each participant since each participant experiences a unique sequence of trials. This allowed us to investigate cortical regions underlying the surprise as

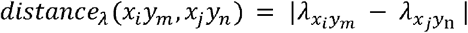

This RDM explained pattern similarity in dorsomedial prefrontal cortex (dmPFC). In other words, this region’s neural patterns were similar when similar surprising changes were registered (Fig 6c, right), and this was true in all three domains.

The above results suggest that, even though there was no explicit need to predict nor a sequential pattern, in the current learning tasks predictive dynamics were correlated with dmPFC neural representations. This highlights this region’s independence from task state and task coordinate computations in the vmPFC and amPFC respectively.

We also assessed collinearity between the three model-derived process variables *φ*_*t*_, *λ*_*t*_, *δ*_*t*_ by correlating them within participants. Correlations were small across domains. One-sample t-tests were performed, comparing correlations against zero after Fisher transformation; these were only significant for the correlation. () Spatial (Mean = −0.017, p = 0.20), Social (Mean = 0.018, p = 0.138), Sequential (Mean =0.005, p = 0.66) () Spatial (Mean = 0.069, p < 0.0001), Social (Mean =0.10, p < 0.0001), Sequential (Mean = 0.077, p < 0.0001), () Spatial (Mean = 0.005, p = 0.405), Social (Mean = 0.008, p = 0.66), Sequential (Mean = 0.019, p = 0.26). Overall, these weak correlations, suggest that these regressors capture distinct computations (Supplementary Fig. 10).

The RSA results so far outline a novel and important finding. The prefrontal neural patterns did not show specificity to features in particular domains of experience but rather to implementing different domain-general computations. Each PFC region reflected distinct computations for estimating states, coordinates and dynamics, each of which may be important for building full internal model.

### Prefrontal Computations are Invariant Across Learning and Inference

The results above show that during learning of internal models, the prefrontal activity patterns reflect distinct computations, that are engaged in similar ways across domains. This suggests that each region is specialized for a distinct computation; specifically, changes to task state representations in the vmPFC, reference frame representations in the amPFC and predictive surprise representations in the dmPFC.

It should follow that, during the test phase participants will infer the hidden feature based on these learned internal models (Fig 3a, Fig7a Trials), using similar computations^14,19^. If this is the case, it would mean that the computational demand on the midline PFC during the test phase, could be similar to learning computations which are invariant across domains. However, because the test phase places different demands on inference and choice, we assumed that the participants may recruit these computations in an adapted form appropriate for test trials.

During learning, participants were instructed to make an overt categorization based on the x-feature. Nevertheless, we hypothesized that they would also (implicitly) learn the latent task state by integrating the z-feature, updating beliefs about which hidden state (x_i_y_i_z) generated the observations even though this was not required by the explicit instructions. In the test phase, the participants needed to make a judgement based on the z-feature value: whether the hidden y feature is y_1_ or y_2_. For this, they needed to infer which of the two Gaussians was more likely to have generated the z-feature values presented sequentially in each test trial. We assumed equal prior probabilities for choosing a state (p(s_1_) =p(s_2_) =0.5) and quantified participants’ belief based on updating this based on the relative likelihoods of the two exemplars under each category (see Methods). This results in different predictions for each type of test trial. EE trials would entail a strong prediction favouring one category. A1A2 have high uncertainty (since the evidence is balanced, and there is no correct answer). We submitted these values for each trial as a parametric modulator in a GLM fitted to the inference task. In the subsequent analyses in this section, effects for each participant were averaged across the domains, since PFC regions previously showed no evidence of variation between domains. Following averaging, participant maps were entered into a group analysis testing for effects of the modulator, which were thresholded at q⍰<⍰0.05 (FDR corrected) with an extent threshold of 25 voxels.

We found a cluster of voxels in vmPFC which showed activation that was predicted by the strength of belief in a specific state (Fig 7b Top). That is, vmPFC activity indexed the relative evidence for a particular state during the test phase.

This prompted us to test similar evidence for amPFC and dmPFC as follows. As described above, the posterior distribution of the winning category shifts its location during test trials. This can result in a parametric modulator that predicts higher BOLD activity with large changes (e.g., 180 degree) in posterior update direction (between test trial stimuli 1 and 2), compared to none (0 degree). This is analogous to *φ*_*t*_ used during Learning. In other words, as participants update their beliefs about the state going from exemplar 1 to exemplar 2 this yields a frame of reference in which the direction of the update is computed. Consistent with this, we find directional shifts are correlated with amPFC activity but not the other PFC regions (Fig 7b Middle). This suggests that trials where participants had to shift frames generated more amPFC activation than trials which did not require this, denoting its importance in maintaining a reference frame.

dmPFC representations during Learning were sensitive to predictive surprise. Specifically, the *λ*_*t*_ metric we used quantified the trial-by-trial by computing the likelihood of next observed z-feature value under the previous posterior. We reasoned that during test phase, a similar computation would predict higher BOLD response in this region owing to change in task policy. Based on the posterior obtained after observing the first test exemplar, the likelihood of second test exemplar could be then computed. If the z-feature value observed in the second test exemplar has a lower probability under its predictive likelihood, the more surprising it would be. And this could result in a higher BOLD activity. This modulator showed an effect in dmPFC; activation was higher when participants experienced more surprising transitions within a single test trial (Fig 7b below).

These results are compatible with the idea that learning and inference recruit similar processes within the PFC. In other words, a same set of specialized computations used for searching over the parameter space of latent world models is be reused when making inferences based on this world model, in line with a flexible cross domain model-based inference and decision circuitry. We interpret this to suggest that the process of building an internal world model, underpinned by the same three computations, which was invariant across distinct looking domains and recruited similarly in learning and test phases. Our results provide to our knowledge, the first evidence of such a principled computational scheme organized for world modelling in the PFC.

## Discussion

The human medial PFC is known to learn flexible and generalizable internal models. We set out to test whether the functional specializations that enable the PFC to achieve this were domain-specific or computational. To this end, we developed a novel design where learning occurred across different domains (spatial, social, and sequential) with distinct surface features but where all domains were governed by the same latent task structure. The domain-specific features were designed to be as distinct as possible to test whether PFC neural patterns were sensitive to specific domains of experience. However, the observed features of the stimuli failed to explain PFC activity patterns. We therefore asked whether PFC activity instead reflects operations on internal models used for learning and inference. We found we could capture participants’ behaviour in a test phase with a Monte Carlo sampling model that was specific to each participant. The model casts learning as mapping observed features into a latent state space. Neural patterns in vmPFC were predicted by changes to the location within this latent space. The amPFC patterns were tuned to local directional shifts within this latent space for each state and switches between states (i.e., latent spaces) in what appeared to be a global task coordinate system. This coordinate system organized different states orthogonally. Finally, the dmPFC patterns reflected prediction of upcoming observations and signalled task policies. In all three cases, the functional specialization was consistent across the three representational domains. Importantly, the same functional specializations were also evident during the test phase, suggesting these computations generalize beyond learning to model-based inference. These results suggest that world models in the midline PFC are not organised by domain of experience. Instead, the functional specialisation here reflects principled biases for suited for different types of world model computations.

Learning, tracking and inferring the state of the environment is the core computation we have probed in this study. A state here is the identity of the Gaussian that generated the observed stimulus with its x, y and z features. Learning the state parameters in the learning phase is central to later inference, where the y feature is hidden. We assumed that each state’s expectation of its μ parameter is a point in the z-feature latent space; these points change their locations from trial to trial until learning converges on two locations, one for each y feature (i.e., the mean of each Gaussian distribution). vmPFC activity patterns were predicted by location changes in this latent space. Importantly, vmPFC activity was predicted by the latent state parameters but not by the observable features of the stimuli themselves. This suggests that vmPFC had an abstract internal model of the task, encoded in a unidimensional space, consistent with the dimensionality reduction observed in this region during other learning tasks^15,16^. During learning, this region appeared to perform inference over parameters within each state, tracking the change, while in test phase it performed inference over states themselves. This suggests model-based probabilistic inference adapted to task as a core computation of this region. Many studies of vmPFC emphasise its role in coding the reward value of stimuli^29,30^. However, in our learning phase, acquisition of the critical information was implicit, with no explicit reward, feedback or value cue guiding participants. This implies that the core function of vmPFC may be to flexibly acquire and update internal models and not to explicitly code rewards. This might also explain why non-task related stimuli might be encoded here, even when not rewarded, if they form part of the underlying model^28^.

Another key interpretation of vmPFC is that it encodes a cognitive map of task variables^7,14,32^. Our results extend this interpretation in two ways. First, the process by which vmPFC acquires a map is not presently clear. Approximate Bayesian inference appears to be a plausible candidate based on our findings. Secondly, instead of a static deterministic map, it could be that vmPFC encodes a fully searchable graph, a claim which would be consistent with reinterpretations of hippocampal codes for cognitive maps posited recently^65^. Our modelling approach used a Monte Carlo sampler to do approximate Bayesian inference. While this is not the only model that may be compatible with our data, it does provide an account of vmPFC that is consistent with other findings. If the vmPFC works as a kind of sampler, then we would expect lesions to this region should result in high variability in state-based decisions, due to high variability during sequential drawing of samples. This is what a recent neuropsychology work on vmPFC lesioned has humans demonstrated^66^. More generally, we suggest that probabilistic learning and inference of task states could be the singular computation supported by vmPFC, offering a parsimonious account across studies.

The second computation we investigated was direction shifts in the latent space. Many states of the world need to be traversed for various goals. In our task, four different states are learned (x_1_y_1_z, x_1_y_2_z, x_2_y_1_z, x_2_y_2_z) and participants move regularly between these states as they view different stimuli. In real-life situations, it is important to keep track of direction change in a state. For instance, is the belief about a person (or object) changing consistently in the same direction along an axis (e.g., increasing professional conduct or size of object) or did a sudden direction change occur (e.g., rapid increase in romantic behaviour or decrease in object size). Thus, a local reference frame for each state has to be computed. Similarly, as states switch, a global reference frame across states needs to be encoded as well. We found that amPFC encodes both of these, suggesting a global task coordinate system. Our model assumes that the four task states were orthogonally separated from each other. This 2-dimensional organisation has been proposed in social inference tasks when participants have to encode people along two distinct social dimensions^19^. Here we show that such computations are not limited to social inference. At first, this may seem odd with studies demonstrating amPFC during social updates and inference. We would argue that it is the reference frame computation rather than the feature representation that amPFC brings to the table.

Social inference requires adaptive changes to one’s reference frame^17–19^ (say about status, mental states etc) of another person. Such frame switching is important in real-life situations when considering and switching between different goals in the same set of states^26^. Thus, our data suggest a novel way in which humans organize their internal models. By representing goals as directions within a frame in amPFC, we may be able to represent multiple simultaneous states while avoiding interference between them. Task coordinate computation has been shown to be important^24,26,67^, and is absent in most contemporary frameworks of model-based inference. This computation is also needed during tasks requiring relational problem solving using a frame. For instance, understanding the instruction ‘it is behind the door to my left’ requires constructing another person’s reference frame, translating their coordinates into one’s own, and maintaining both simultaneously. Thus, a relational computation that depends on the frame itself, not just the agents (or objects) within it. amPFC engagement is observed consistently in such tasks^27^, across studies that seem superficially different but represent a conceptually similar problem. We thus suggest that constructing and maintaining the current task coordinates is a generic computation performed by amPFC.

The final element of world models that we investigated is forming task policies. The first two computations helped in inferring and representing the world in efficient formats. The final computation allows people to make temporally abstracted actions using the model. For this we assumed that participants had a forward predictive model, representing future task representations, with requisite task policies. These was monitored actively resulting in keeping or changing them based on the validity of the predictions. This was formalized as the surprise, the inverse of the likelihood of observing the current trial given the previous one. Crucially, we found that participants’ response times were modulated by the surprise encountered in a trial, independent of the perceptual features of the exemplar. Across domains, we also found that dmPFC representations were predicted by how surprising participants found each trial, given their forward predictions. In learning, this suggests that even though the overt categorization was independent of the internal model dynamics, the policy instantiated in the dmPFC did influence this downstream decision, as reflected by increased response time. During test phase, different conditions elicited distinct surprise values in second test exemplar, given the first, which also modulated their response times. This likely resulted from differences in whether participants stayed in the predicted state with a different policy (press left if s_1_) or switched to a new state with a different rule (press right if s_2_). This suggests that dmPFC does not merely register surprise, but tracks the computational cost of maintaining a policy given the unfolding trajectory.

dmPFC has been implicated mainly in action tasks^21^, strategy formation^33^ and planning^37^ in general; all of these require finding optimal ways to take actions across task states. This is also what was observed in our recent study^23^ when during naturalistic experiences, activity increases in this region were observed when participants’ beliefs about expected future trajectories was changed, due to surprising new actions altering what was about to unfold. A task policy (or strategy) is a specific set of such actions across states chosen by participants to make decisions, and we suspect this computation is what the dmPFC performs. This generic computation can also explain why dmPFC is implicated in tasks seemingly unrelated to one another. For example, offline exploration tasks such as mind wandering^35^ have been shown to recruit this region, as well as observing others’ actions. In the first case, mind wandering may involve exploring different policies (kinds of action sequences) across states in one’s internal model, even in the absence of an explicit a goal. In the second, action observation may trigger transitions between states in the internal model of another agent, offering a way to infer their policies. Similarly, the common occurrence of dmPFC activity in (unsigned) prediction error^36^ – which is roughly, the magnitude of surprise - across tasks can be parsimoniously explained by online shaping of task policies for better responses. In other words, this region may not be encoding the prediction error as much as forming adaptive representations for task policies. These require inference by searching a specified space in unique paths. Just as motor regions maintain multiple concrete action plans^68^, the dmPFC may maintain multiple abstract task policies over states, switching them adaptively by their predictive validity. On this view, dmPFC is most engaged when a task behaviour requires multiple distinct ways of acting across different latent task states and revising that search online when new evidence violates predicted outcomes.

We end by considering some implications of the Monte Carlo sampling approach we used to model learning. Our model assumes that PFC may be generically inferring latent environment structure using probabilistic inference. Crucially, however, we did not assume that this extraction is optimal, given the resource constraints the human brain has. Sampling^47,48,69^ offers a computationally feasible candidate approach to implementing approximately Bayesian inference about an open space of possible world models in the cortex. Rather than weighting an infinite set of possibilities explicitly, brains approximate this implicitly by forming a generative model, from which an arbitrary number of samples can be drawn as needed. Moreover, sampling representations can naturally explain neural variability and behavioural stochasticity, which otherwise must be explained away. Sampling only becomes exact in the limit of infinite samples while online inference trade-offs between the costs and rewards of additional computation. Our Monte Carlo model embodies this idea. It is inspired by resource rational (computationally bounded) models that have explained a wealth of behaviour in higher-order cognition^44–46,59,60,70,71^. In our task, participants were sensitive to the sequence of test trials presented. This result is incompatible with normative analysis of the task. We took this to suggest that participants might be reusing the samples previously used. The most striking feature of the behavioural data was a consistent within-subject bias and order effects. We found we could fit this with a model which assumes participants approximate the likelihood by a comparison process between internally simulated data and external observation, where a resourceful reuse can exhibit bias. Such schemes have important theoretical implications in machine learning and might be useful to explore in cortical networks^72^.

In closing, we suggest that humans learn and reason over internal world models using three different regions in the PFC, specialized for distinct computations. Together these computations enable probabilistic discovery of task states, organizing multiple such states into a coordinate system and forming task policies in ways that are invariant to general task demands and are similar across domains with distinct features. Such a computational scheme could potentially unify a wealth of PFC results across different studies using seemingly distinct tasks and stimuli, into a singular role as the brain’s general-purpose model-based reasoning system.

## Methods

### Participants

The study included 31 young adult participants (MeanAge=22.84, SDAge = 3.9, range 19-37, 81% female). All participants were fluent in English, and had no history of neurological or psychiatric illness. The study received ethical approval from the Edinburgh Psychology Research Ethics Committee, and informed consent was collected from all participants. A screening was also conducted beforehand using a very brief categorization task online using simplistic shapes to determine their cognitive aptitude. Only those participants who had more than 50% accuracy in this were deemed eligible for the main study.

### Stimuli and Materials

Participants completed three learning tasks within three virtual worlds that represented distinct domains of learning: spatial, social and sequential. Within each domain, after the learning task they were immediately administered a test phase before the next domain. Domain orders were counterbalanced across participants.

A set of images and video stimuli for the three worlds were generated using the video game Age of Empires II (Definitive Edition). Using its in-built map editor, we generated virtual world stimuli for each domain and category (see Fig 1A). Each domain had 16 distinct exemplars each with a categorical feature x, a categorical feature y and a continuous feature z, which was associated with the categorical feature y. For example, in the spatial domain the x feature was whether the village had gold (x_1_) or stone (x_2_) mines. The number of mines, z, was unrelated to this category but could be used to predict whether the scene included a tent (y_1_) or tower (y_2_). The feature z was drawn from two Gaussians (σ = 3) with two possible means (μ = 7 or 14; shown in Fig 1C). Exemplars containing y1 had the z feature drawn from the lower Gaussian and those containing y2 from the higher Gaussian. X and y features were fully crossed, for four stimulus types: x_1_y_1_, x_1_y_2_, x_2_y_1_ and x_2_y_2_. To set z, we drew 4 exemplars from each of these combinations using the 5^th^, 25^th^, 75^th^ and 95^th^ percentiles of the respective Gaussian to produce 16 unique stimuli. For example, x_1_y_1_z_1_, x_1_y_1_z_2_, x_1_y_1_z_3_ and x_1_y_1_z_4_ denotes four exemplars from x1y1 category of exemplars with different number of gold mines, drawn from the Gaussian *N* (7,3).

This feature triplet mapping (x,y,z) and, importantly, the same Gaussian latent structure was used in other domains as well, making the task structure consistent across them. In the social domain, exemplar videos of two clans of bandit queens appeared (blue/red clothing as x feature). These queens could engage in a professional or romantic manner with a local town mayor. The mayor behaves in a distinct manner in each case; offering silk gifts (y_1_) or trading weapons (y_2_). The speed with which the queen moves was determined by the z feature, drawn from the two Gaussians, thus linking her speed to her mental states.

The sequential domain had the same underlying structure. Exemplar videos showed an object (lumber pile) moving across space while participants categorized it as requiring actions of cutting (x_1_) or burning (x_2_) it based on its colour. The object was transported in specific sequences across the screen, first vertically and then horizontally, ending up on the left (y_1_) or right (y_2_) of the screen. Here the length of the vertical movement (z) was drawn from the two Gaussian distributions.

All exemplar images and videos used during learning and inference were approximately 1920 x 1080p resolution and were cropped and aligned using grids overlaid within the in-game editor engine.

### Task Design

There were three phases to each task; i) a pre-training, ii) learning (training phase) and iii) inference (test phase). Only the latter two stages were completed within the scanner. Each domain had its learning and then inference completed in a single scanning run, before moving to the next domain. The order of domains was randomized across participants, and it was the same order as they experienced in the pretraining phase (see below).

### Cover Stories

Each domain came with its own cover story to induce a richer world-building experience for the participant. In spatial domain, the cover story was that each scene represents a village that is controlled by either of two families (Pine/Winds). The classification task was to identify the family owning each village (determined by whether its mines contained gold or silver). At the same time, they were instructed to learn the relationship between each scene’s number of mines and the associated building that is observed (Tent/Tower). During the test phase, they were told that a wizard cast a shadow over the building and they must infer its identity (Tent/Tower) from two scenes. In the social domain, a mayor of a town is in talks with local bandit queens from two clans (Phoenix/Wolf). The mayor can provide the queen with weapons (Professional) or give her a gift of silks (Romantic). Participants must categorize the queen’s clan (determined by the colour of her clothing) while learning about how her behaviour predicts the mayor’s gift. During the test phase, participants were told the mayor’s wife now knows about his infidelity. The mayor is no longer able to express his feelings through gift-giving so participants must infer his feelings from the bandit queen’s behaviour. Finally, sequential learning involved a cover story of participants running their lumber business by ‘acting’ on each exemplar (pile of lumber). The categorisation task was whether the lumber should be Cut or Burned (determined by the colour of a flag placed on it). This was in addition to learning whether the lumber would finish on the Left or Right side of the screen, based on how much it moved vertically. The test phase only showed the vertical movement, and participants had to choose whether it should be moved to left or right.

### Pretraining

In the pretraining phase, participants were given high-level instructions about the three tasks outside the scanner. For each domain, they were shown only the x and y feature together as exemplars. In this phase, the z feature only had 1 mine in spatial domain while a static image was used in social and sequential domain (i.e., both the bandit queen and lumber did not move). In each domain, participants learned the categorizing rule from these exemplars via trial and error. This rule was unique to each domain and participants learned it until they were confident and had <10% error rate on pretraining trials. They were then shown dummy trials of how Learning phase would occur inside the scanner and told that they would be tested on the association of the y and z features, later during the test phase. For example, in the spatial domain they were told the number of mines would vary and that later they would be asked to predict the correct building (tent/tower) with only the mines to be used for inference, implying a relationship between them. Similar logic was used in the other domains. They were also shown a dummy trial of how the test phase would appear. They were reminded of these instructions in the scanner before the test phase began by visually presenting the same instructions again for 30 seconds.

### Learning

Learning phase was conducted in the MRI scanner and started after participants were well accustomed to response key control. There were brief instructions to recap for 30s before each task started. During learning, participants were instructed to apply the same binary categorization rule they learned in each domain (x_1_/x_2_). Each learning trial was of 2.5s duration for spatial and sequential domains, and 3s for the social domain. This was followed by an interstimulus interval (ISI) jittered between 2 and 3s. A feedback image (2s for spatial and sequential, and 1.5s for social) appeared afterwards depicting whether their categorization was correct or not. Importantly, participants were also attempting to learn the association between y and z but they made no overt responses on this basis and therefore received no feedback whatsoever for this aspect of learning. Trials were separated by a blank screen with fixation (ITI) of duration jittered between 2.5s and 3.5s. Jittered durations were used to optimize for better neural pattern estimations.

Each learning task occurred across three blocks, separated by rest blocks of 25s where a blank screen was displayed. A pre-Learning rest block of 25s was also administered before the first learning block started. At the end of each rest block, a 2s image notified the participant that the learning block was about to start. Each category state (x_1_y_1_z, x_1_y_2_z, x_2_y_1_z, x_2_y_2_z) had 4 unique exemplars (see Stimuli above). All z_1_ and z_4_ exemplars were shown once only in a block while z_2_ and z_3_ exemplars were shown twice (following the underlying Gaussian distribution). This resulted in 6 presentations of each x_i_y_i_ state, for a total of 24 trials per learning block and 72 trials per domain. Post learning, they read the instructions for the test phase for 30 seconds before this commenced.

### Test Phase

The test phase was administered immediately after learning. During the test phase, each trial consisted of two sequentially presented exemplars (Fig 3a), where the y feature was hidden and z feature took new values not presented during learning. Participants were told that both exemplars within a trial shared the same hidden y feature and were asked to infer this feature. Correct inference required using the learned mapping from z to y, that is, determining which of the two Gaussian distributions was most likely to have generated the observed z values. In the spatial domain, the building (y) was hidden (shadowed out) and only the mines were visible. During the social domain test phase, the mayor did not offer goods and thus his intent had to be inferred from the movement of the bandit queen. Similarly, in the sequential domain, the lumber only moved vertically down and where it would end up (left or right) had to be inferred by the participants.

In the spatial domain, trials involved presentation of two back-to-back images of each 2s duration. Social and sequential domains involved two videos of 2.75s and 2s respectively. Participants could only respond during the second exemplar on each trial. An intertrial interval jittered between 2.5-3.5s was interspersed between trials. There was no feedback. Inference was conducted over three blocks, with each test block having 24 trials with a 5 second rest block in between, for a total of 72 trials.

We devised 7 distinct conditions for trials in the test phase. Test exemplars were sampled from three different regions of the two Gaussians (Fig 3b) and could be presented in different orders, yielding seven test conditions. Some exemplars were taken from the extreme ends of the two gaussian distributions (labelled E1, E2), which are only associated with one of the two y features. Other exemplars were taken from the central region in which the two gaussians overlapped. These exemplars predicted both y features with similar probability so they are labelled A1, A2 (for ambiguous). By pairing extreme and ambiguous exemplars, we are able to vary the difficulty of inference in the test phase. For example, on EE trials, both exemplars were extremes (E1E2), providing strong evidence for the correct y feature and serving as an easy way to confirm if participants had learned the task. All EE exemplars were novel (i.e., not presented during learning), ensuring that test performance required generalization to new values and there were 18 such trials.

A second set of trials (AA) featured two ambiguous exemplars. Because their z values lay in the overlap region, there was no objectively correct y feature. For presentation of results, we arbitrarily coded responses choosing the y_1_ model as correct. AA trials were presented in both possible orders (A1A2, A2A1), with 9 trials per order (18 trials total). The remaining trials were mixed extreme– ambiguous (EA/AE) trials, in which one exemplar was extreme and the other ambiguous. We created four scenarios: E2A1 and A1E2, where the ambiguous exemplar was closer to the correct Gaussian, and E2A2 and A2E2, where the ambiguous exemplar was closer to the incorrect Gaussian. Each of these four scenarios was presented on 9 trials, yielding 36 EA/AE trials. In total, the test phase comprised 72 trials (18 EE, 18 AA, 36 EA/AE). This test structure design allowed us to systematically vary the strength and ambiguity of the evidence used for inference, as well as the order in which exemplars were observed, thereby distinguishing probabilistic inference from heuristic or sequence-insensitive strategies.

After test phase, participants were debriefed and asked what strategies did they choose for each domain from a candidate set of strategies (data not analyzed in this study).

### Neuroimaging Data

Functional MRI data were acquired on a 3 T Siemens Prisma scanner (32-channel head coil). Each participant completed one whole-brain multi-echo EPI run for each domain (46 slices covering the whole-brain; TR = 1.7 s; TE = 13/31/50 ms; flip angle = 73°; 80 × 80 matrix; 3 mm isotropic voxels, slice thickness = 3.0 mm and multiband factor = 2). There were 688 volumes for the spatial domain with social and sequential domains having 788 and 729 volumes respectively. A high-resolution T1-weighted scan was obtained with an MP-RAGE sequence (1 mm isotropic; TR = 2.5 s; TE ≈ 4.5 ms; flip angle = 7°).

### Pre-processing

Functional images were pre-processed and analysed using SPM12 and the TE-Dependent Analysis Toolbox (Tedana)^74^. The first 4 volumes of each run were discarded. Estimates of head motion were obtained using the first BOLD echo series. Slice-timing correction was carried out and images were then realigned using the previously obtained motion estimates. We used Tedana to combine the three-echo series into a single time series and to perform multi-echo independent-component analysis (ME-ICA). This divided components of the data into either BOLD-signal or noise-related. The noise components of the data were discarded. This process reduces motion effects and susceptibility artefacts in ventral and orbitofrontal cortex^73,74^. Then images were unwarped with a B0 fieldmap to correct for irregularities in the scanner’s magnetic field. Finally, functional images were normalised to MNI space using DARTEL. The spatially normalized images were smoothed with an 8 mm FWHM Gaussian kernel. Smoothed images were only used in the parametric modulator GLMs (Fig 7b). Unsmoothed images were used for the multivoxel pattern analyses.

### General Linear Model

After preprocessing, for each domain, we separately built two sets of general linear models (GLM) to generate single trial estimates for multivoxel pattern analyses. Each modelled the BOLD data with a high-pass filter with a cut-off of 180 sec. We used the six motion parameters and their first order derivatives as covariates in all GLMs. These two GLMs were designed for the two kinds of searchlight RSAs we performed and the logic is as follows. For the Stimulus-based RSAs (Fig 2), we pooled learning exemplars over blocks to obtain a single pattern estimate for each exemplar. This was because the stimulus values used for RDMs were same in all blocks. In contrast, in the Process-based RSAs (Fig 6), derived from the Monte Carlo sampler fitted to each participant, each individual trial had a unique value and thus trials from the three learning blocks had to be modelled separately.

For the Process-RSA, for each domain, we ran a single GLM on the unsmoothed BOLD time-series to provide single-trial parameter estimates. Some exemplars were presented twice in each learning block. These were modelled with two separate regressors for the first and second presentations. Thus, there were 72 learning regressors in total. Additional regressors modelled the Test phase trials (as a single test regressor), feedback presentation, the three rest blocks (separately) and two response time duration regressors for learning and inference trials each. Thus, each GLM had a total of 72 main learning-trial regressors, and 7 ‘other’ nuisance regressors. For Stimulus-RSAs, we used the same approach but used a single regressor to model the three presentations of each exemplar over learning blocks. Thus, this model had 24 stimulus regressors rather than 72.

We used univariate noise normalization, i.e. the t-maps estimated from these GLMs as the input for the RSA.

### Wholebrain Searchlight RSA

We used searchlight analyses in CoSMoMVPA^75^ to investigate how activation patterns were predicted by various theoretically motivated RDMs. We submitted the neural patterns obtained from the GLMs above to a wholebrain searchlight RSA. We used a spherical searchlight with radius of four voxels. We computed pairwise distances between neural activation patterns for each exemplar using the squared Euclidean distance. Spearman correlation was used to assess similarity between theoretical RDMs and neural RDMs, in which we also removed the mean pattern. In Stimulus-RSA, RDMs of dimension 24x24 were compared between neural and theoretical levels to obtain the correlation value for the centre voxel of each searchlight. In Process-RSA, this was obtained separately for each learning block (again with 24x24 RDMs), and the final correlation values obtained by averaging across all three blocks, for each participant in a domain. Correlations were Fisher-z transformed for group-level analysis to map the correlation values to an approximately normal distribution.

To determine which voxels showed statistically significant correlations between neural and theoretical RDMs, we used the Monte Carlo function within CoSMoMVPA to run 1000 simulations at the group-level (by sign-flipping across participants). This resulted in a set of p-values corrected for multiple comparisons with threshold-free cluster correction (TFCE). We used a threshold of p<0.01 and only showed the voxels exceeding this value. Whole-brain maps are visualized as t-maps.

The structure of the RDMs used in these analyses are reported in the Results section. In the RDMs, during the first trial where there is no update to any of the measured metric occurring, we put a value of 0.

### Behavioural Data Analyses

Participants’ categorization accuracies in Learning were computed for each domain, averaged across all exemplars. Test phase accuracies were computed for each scenario and averaged across all exemplars. AA trials had no correct answer so y1 was arbitrarily classed as the correct response on these trials, so they could be visualised alongside the other conditions. Thus, we used seven scenarios – E1E2, A1A2, A2A1, E2A1, E2A2, A2E2 and A1E2. We removed participants with less than 50% accuracy on all except AA trials (as there is no right response on these trials). This resulted in exclusion of 2 participants from spatial and social domains, and 5 participants from sequential domain.

To assess if surprise measured modulated the response time, we used a mixed-effects model pooling across domains, separately for learning and test phase. Here the dependent variable was response times per trial (removing non response trials) with the associated surprise as the independent predictor. Random effects over participants and domains were also included.

### Learning Models

We compared the following models of learning, all fitted to participants’ test phase behavioural data (in which decisions are based following the presentation of two test stimuli). Models were trained and tested separately for each participant in each domain using leave-one-condition-out-cross-validation (LOOCV), as described in the Results.

### # Random Responding

We put 0.5 as the choice probability for each scenario during model fitting and model recovery for this learner.

### # Boundary Learner

As one alternative model, we considered a simple strategy wherein participants learn a decision bound. Such models have been shown to capture participant behaviour in various categorization tasks^53,54^. The decision boundary learning compares each z feature value in the test data pairs to a boundary value. This is a decision heuristic which instead of encoding learning exemplars and their statistics, just collapses to a boundary value which is used to compare test exemplars as belonging to either category state. Since the standard deviations were equal, the ideal value of this boundary, c is midway between the mean of the two distributions as s_1_

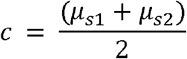

But our boundary learner assumes a noisy encoding or recall, making this boundary differ across participants, and is thus a free parameter. Probability of a state *s*_1_ given the first test data 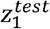 then is

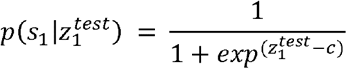

Probability of a state *s*_1_ given both test data would be the product of both pairs

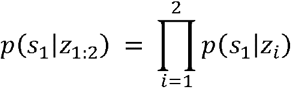

Final decision probability for a state *s*_1_ (compared to *s*_2_) then is computed as

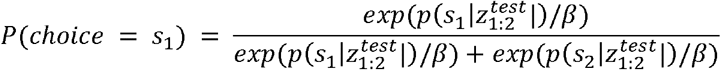

Where, *β* is the temperature parameter which imposes decision noise on the choice probabilities. Participant test behaviour data was fitted through the L-BFGS-B optimization algorithm in R^76^.

### # Prototype Learner

As another alternative model we considered a prototype learner. This model compares features of each exemplar, to a prototype acquired during learning phase. Neural correlates of this type of learning have been shown^55^ and studies have shown that recency weighting of the data explains human data better than equal weighting^56^. The model compares the test z values to category means *μ*_*s*1_ and *μ*_*s*2_ to obtain a distance from each. It weighs this distance with an attentional weight parameter *w*. Since there is only one feature (z values), this parameter differentially weighs the two test data pairs^56^.

This weight of z feature in first exemplar in the pair are defined as

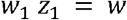

The weight parameter *w* determines how much weight is assigned to first exemplar. And similarly for the second exemplar,

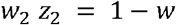

An ideal learner would weight both exemplars equally, but the weight parameter allows for the possibility that participants show a bias towards being more influenced by the first or second exemplar in the pair.

Probability of a category state *s*_1_given both test data would be the sum of the distances of each test value to the model mean

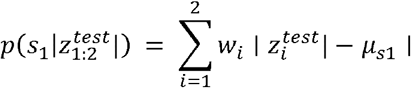

Final decision probability for a state *s*_1_ (compared to *s*_1_) then is computed as

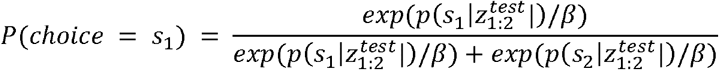

Where, *β* is the temperature parameter which imposes a decision noise on the choice probabilities. Participant test behaviour data was fitted through the L-BFGS-B optimization algorithm in R.

### # Normative Bayesian Learner

#### Learning Phase

To model the ideal Bayesian learner, we computed a posterior distribution on mu and sigma for each category s_1_ and s_2_ conditioned on the training data.

The model assumed the *z* values were drawn from a Gaussian parametrized using unknown *μ* and precision *τ* (*τ* = 1/ *σ*^2^)

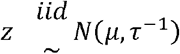

The mean parameter *μ*, conditioned on the precision, is assumed to be drawn from a Gaussian with a mean of 0 and precision of 0.001 (i.e., weak hyperpriors)

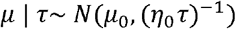

And the precision is assumed to be drawn from a Gamma density

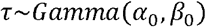

The likelihood of drawing *n* observations of z is the product of Gaussian densities

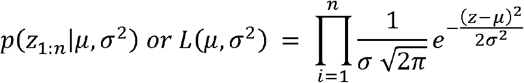

Taking the log likelihood to turn the product into a summation (equation 1)

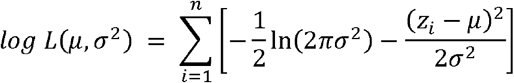

The generative structure above is same for Bayesian learner and its approximation scheme of Monte Carlo sampling, but they deviate in how it is learned.

The Bayesian model learns through conjugacy. The posterior over conditioned on precision upon seeing *z* features for a given state, *s* is

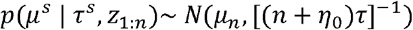

Where, *μ* _*n*_ is the posterior mean parameter 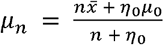, n is the total data points, 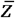 is the (data) sample mean with *μ* _0_ is the hyperprior term for mean and *η*_0_ acting as the prior precision strength.

The Gamma parameters are updated similarly for the precision parameter. The resulting posterior distributions were: x_1_y_1_z/x_2_y_1_z, *N*(µ = 7, σ = 3) and x_1_y_2_z/x_2_y_2_z, *N*(µ = 14, σ = 3).

#### Test Phase

During test phase, the two test data points for each trial are then used to compute posterior probabilities of each model. The likelihood for a given category state s_i_

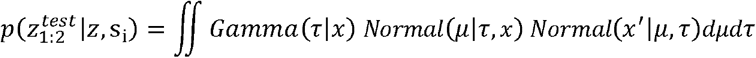

This yields a (shifted) student-t distribution as posterior predictive^77^, which is the Bayesian conjugate for a Gaussian with unknown parameters.

The decision probability for a state s_1_ then is computed as

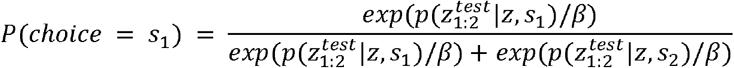

Where, *β* is a temperature parameter capturing noisy selection of the more probable category. Participant test behaviour data was fit to this model through the L-BFGS-B optimization algorithm in R.

### # Monte Carlo Sampler

#### Learning Phase

Instead of conjugacy as in the normative model, a Monte Carlo sampler tracks sufficient statistics to recursively compute the likelihood.

We begin with the log likelihood (equation 1 above), where *n* is the number of observations. This reduces to

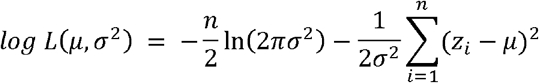

The last term can now be decomposed algebraically to

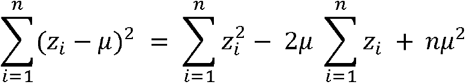

The first and second term here can be replaced by sufficient statistics according to Fisher–Neyman factorization theorem

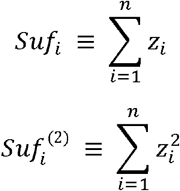

Where *Suf* and *Suf*^(2)^ refers to the two sufficient statistics required to update the likelihood function. The final likelihood at any trial thus becomes

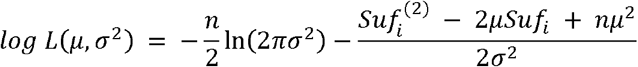

Which can be updated as the data comes in. In other words, one only needs to keep track of the count and sums of the *z* feature values rather than the entire batch.

According to Bayes theorem the posterior is a product of prior and likelihood, normalized by evidence,

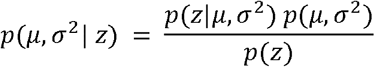

The denominator, p(z) is the normalizing constant, which unfortunately is an intractable quantity in most cases, making the true posterior a hard-to-obtain quantity. Although in simple cases it may be possible obtain the true posterior, we here assume the general case in which it is not.

The goal of sampling/MCMC is to approximate the posterior without ever evaluating this denominator. MCMC exploits the property that we know the posterior is proportional to the product of prior and likelihood up to some unknown normalisation constant

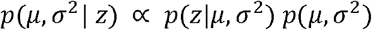

Taking the log to turn this into a summation

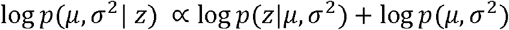

Substituting the likelihood form above and the prior terms, the target (log) posterior density at any trial becomes

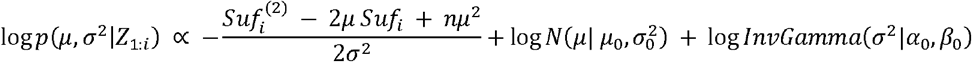

Where the first term on the right is the (joint) likelihood (*Suf* being the sufficient statistics) and the last two terms are the priors on *μ and σ*^2^ respectively.

MCMC draws autocorrelated samples from this distribution. Concretely, at each step, it proposes a candidate *μ*′, *σ*^2′^ using a Gaussian proposal centered on the previous parameter state (*μ* _*t*−1_ if it is the mean parameter) as

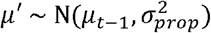

Where 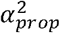 is the width (variance) of the proposal distribution, determining how it explores the parameter space and takes the shape of the true posterior distribution. Similarly, the *σ*^2^ parameter is proposed as

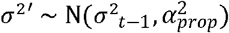

We fixed the proposal distribution width for both parameters to 4 (roughly the variance of the distribution). At each trial the MCMC generated a chain of length 1000. The proposed parameter values were accepted with the Metropolis Hastings acceptance function α as

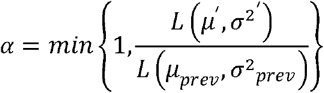

This step, (the Metropolis Hastings ratio) ensures the chain always accepts more likely pairs and stochastically accepts less likely pairs. Asymptotically the chain will visit parameter values with a frequency proportional to their posterior probability. After every learning trial, the sampler estimates the posterior by using the likelihood with updated sufficient statistics. We assume participants carry the samples generated in their final trial for each category over to their test phase inferences.

The spread parameter(variance) is assumed to be drawn from an inverse gamma distribution,

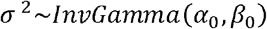

Where *α*_0_= 5 and *β*_0_= 10, again a wide prior relative to the spread of the data.

#### Test Phase

During the test phase, we assume participants use their final set of samples from the training phase to make inferences. Specifically, for a given trial, the probability of the *z*^*test*^ values observed across two exemplars came from one of these y-conditional Gaussians which are now as approximated by a set of samples from their posterior parameters. In other words, what the participants are trying to compute is the quantity,

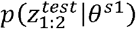

which is the likelihood of the *z*^*test*^ values under the learned parameters of a certain category Gaussians. Comparing this between the categories would result in category posterior probabilities. We assume, however, due to computational resource limitations, participants do not know the exact likelihood function and instead resort to simple approximations of this. This is in line with a range of studies showing humans use a suboptimal number of samples^44^, approximations or a single hypothesis^59,60^ in higher-order cognitive tasks. We formalize this approximation to likelihood using likelihood-free inference, using a framework known as Approximate Bayesian Computation (ABC). Here participant use their learned posterior samples to simulate internal ‘data’, and compare this to the observations.

Under this scheme, the likelihood is not computed exactly, but instead approximated as follows:

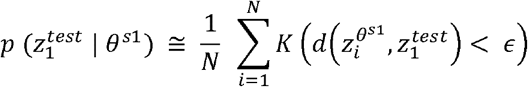

In order to make the process fully distribution and domain-agnostic, rather than analytically computing this, we assume summaries of the posterior are used in place of an analytic likelihood. That is, from the learned samples of *θ*^*s*1^, simulated data 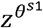 are compared to the actual test data. In our task, we assumed participants compared these simulated data to the observed data using a distance function *d* (we used Euclidean distance) under a tolerance ϵ, which is the only free parameter in the model. We used a grid search where we generated tolerance values from 2 to 6 by increments of 0.1 to fit. A tolerance approaching 0 would mean exact likelihood computation and thus would not be a likelihood-free approximation as it requires a large number of samples to be drawn from the prior, opposing resource rational claims. Importantly, the fraction of accepted samples per drawn samples that survive a tolerance, approximates the likelihood. Under this scheme, the posterior probability for one Gaussian, after seeing the first test value, 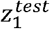 then becomes

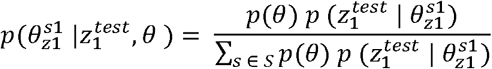

This is the prior probability (0.5) multiplied by the acceptance rates, normalized across the other category Gaussians to return posterior probabilities after first observation. This posterior now becomes the prior for the next and final round of test 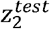. We assume that only the surviving samples are used for the second exemplar in test. Such reuse produces sequential effects based on 1) tolerance used and b) observed z feature order since the likelihood for the second test data 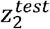 then is now approximated by using samples from the previous posterior (and not the full learned samples) as

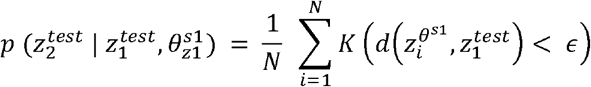

The final posterior over states s_1_ and s_2_ is then:

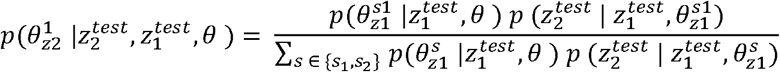

We used these posterior probabilities obtained to form choice probabilities across categories for each participant. The model was fit to the test data with the learned posterior parameters optimizing for the best explaining tolerance for the likelihood-free procedure.

For each participant, we fit both the model and identified the best tolerance value using leave-one-condition-out cross-validation. Importantly, the tolerance parameter used by the participant was kept constant throughout their test phase within a domain.

### Parametric Modulators

For the parametric modulator analysis (Fig 7), we obtained the parametric modulators from test phase results, and created three different GLMs for each modulator, assigning trial-by-trial values to each of the seven conditions within each domain. The key regressor for the test phase was modelled as the onset of the second test exemplar along with its parametric modulator. There were additional regressors for all learning trials, feedback and rest as well. There were three modulators, each analogous to the ones used for creating the learning RDMs. These modulators are as follows (see Results for a more detailed explanation of how the different learning processes were modelled).

For the first modulator, *δ*_*t*_, we assumed participants performed a model selection procedure during each test trial over both exemplars by comparing the total evidence for a specific y feature by computing the likelihood ratio for each Gaussian after observing them sequentially. Specifically, we are interested in the strength of belief (i.e., relative evidence for one state over the other) they had during the second test exemplar 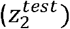 for a state (*θ*^*s*^) after observing the first test exemplar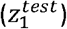, across both states s_1_ and s_2_ (i.e., the y_1_ and y_2_ Gaussians).

The evidence for one Gaussian (y_1_) over the other (y_2_) after seeing the first exemplar 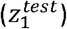 is

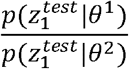

Where *θ*^1^ and *θ*^2^ are the learned parameters of the first and second Gaussian respectively. Subsequently, the evidence after seeing the second exemplar 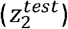 is

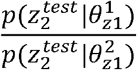

Where 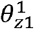 and 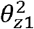 signify the updated parameters of the Gaussian (μ, σ^2^) after observing the first exemplar. To obtain the total evidence we compute the joint likelihood ratio across both exemplars

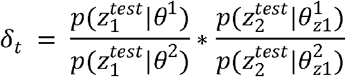

These values were submitted into the GLM as a trial-by-trial parametric modulator of the first computation during test phase.

This approach adheres to the principle of sequential Bayesian updating, where the total evidence for one Gaussian over another is the product of the likelihood ratios from each observation. Specifically, the first term captures the evidence from first exemplar. The second term captures the evidence from the second, given the parameter updates necessitated by the first exemplar. Formally, this represents the joint Bayes factor comparing the two latent states.

The sequential update of the Gaussian parameters within a trial meant that the location of the posterior mean would change from first (*θ*^1^) to second test exemplar 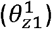. We use the direction change measure we used during the learning phase to quantify this change within a test trial as the parametric modulator of the second computation.

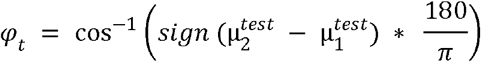

For example, consider the situation of how the chosen Gaussian’s mean parameter changes within a test trial. Take one case, the chosen Gaussian (y_1_) had learned mean parameter 7 and a participant completed a test trial where the first exemplar is of z-feature = 1 and second exemplar is z-feature = 4 (EE condition). Suppose the Gaussian mean after these updates would be ∼4, a decrease in mean. Compare this to another example of the same condition (EE) where the z-features are 20 and 17 respectively. Here, the chosen Gaussian (y_2_), had a learned mean of 14, and upon seeing a data would update to ∼17, an increase. These two updates are in opposing directions where one would be 0-degree and other would be 180-degree.

Finally, after observing the first test exemplar, participants might predict the second test exemplar under both states. We quantified the surprise as how unexpected the second test exemplar was given the first, signalling the surprise.

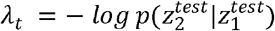

This can be expanded as the posterior predictive, exactly as used in learning, where the likelihood of second exemplar under the state parameters (summed over both states), after seeing the first exemplar. We used the negative logarithm of this quantity to measure the predictive surprise of this transition from first to the second exemplar, under the learned states. If the second exemplar had lower likelihood it would mean it elicits a higher surprise, and thus higher BOLD activity.

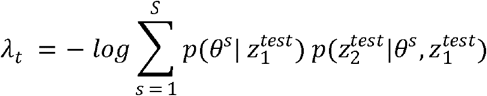

Having calculated these values for each participant on each test trial, we entered them into separate GLMs for each domain. For group analysis, we averaged the maps over domains and thresholded at q□<□0.05 (FDR corrected) with an extent threshold of 25 voxels.

## Acknowledgements & Funding

We’d like to thank Chris Summerfield, Joszef Fiser, Adam Koblinger, Adam Sanborn, Nikolaus Kriegeskorte, and Eric Schulz for helpful discussions. The research was supported by a BBSRC grant (BB/T004444/1). For the purpose of open access, the authors have applied a Creative Commons Attribution (CC BY) licence to any Author Accepted Manuscript version arising from this submission.

## Ethics declarations

The authors declare no competing interests.

