## Supplementary for "Specialized Computations for Generalized World Modelling in Medial Prefrontal Cortex"

Supplementary Materials

### Alternative RDMs

In addition to the stimulus-derived RDMs used in Main Figure 2, we also investigated two other possible mechanisms for representations in the PFC.

Alternative representation 1: Reaction Time

Here, we used the participant's reaction time values for each exemplar trial during the categorization rule during learning. The logic being, since they are categorizing while encoding the latent structure changes, there could be cognitive load related effects, which would be reflected in their response times, and consequently in the neural activity patterns. The RDM is a 24x24 matrix with Euclidean distance between the RT values.

$${distance}_{\text{RT}}\left( i,j \right)=\left| \text{RT}i-\text{RT}j \right|$$

On trials where participants failed to respond, we set the RT to the maximum value observed for that domain.

Alternative representation 2: Difficulty-based RDM

Here, we assumed that exemplars where z is drawn from the area where both gaussians overlap would be more difficult to process than exemplars where z is drawn from the outer extremes of the Gaussians. This RDM coded exemplars according to similarity in difficulty. The resulting binary RDM is defined as

$${distance}_{difficulty}\left( i,j \right)=\begin{aligned} 0,&\text{if }\left( x_{i}y_{m}z_{r} \right)_{\text{ and }}\left( x_{j}y_{n}z_{s} \right)_{\text{ with }}\left( r,s \right)\in\{1,2\}\times\{1,2\}\text{ or }\{3,4\}\times\{3,4\} \\ 1,&\text{otherwise} \end{aligned}$$

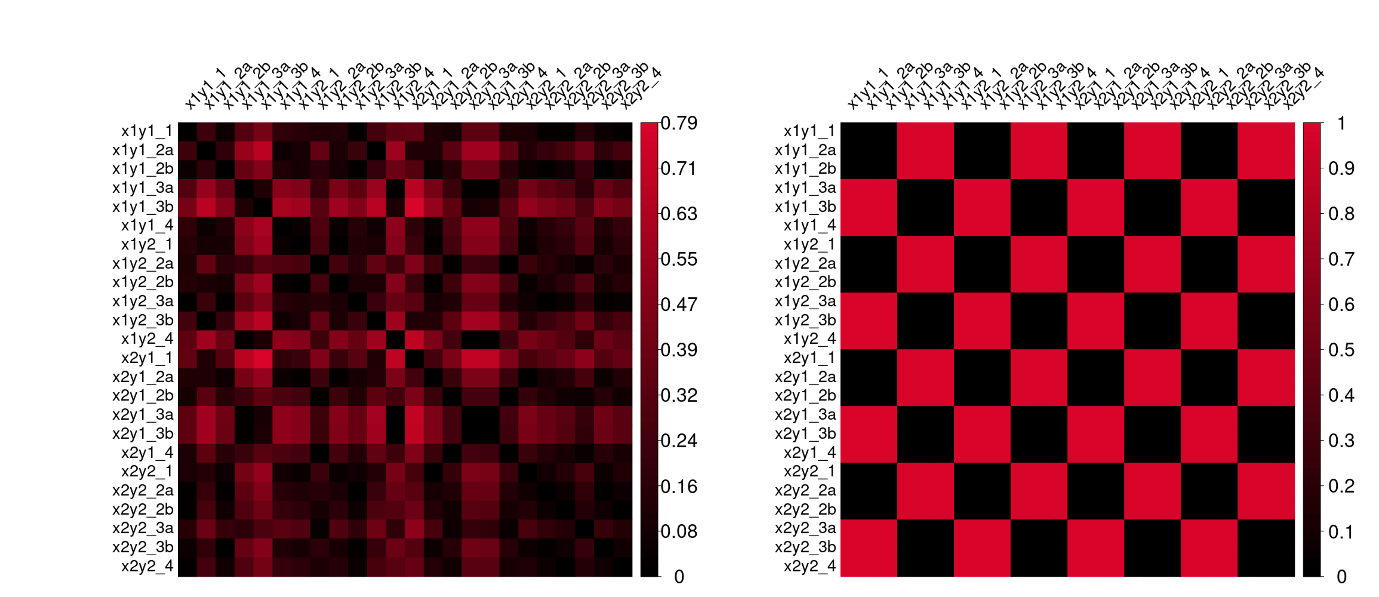

**Supplementary Fig 1: Alternative RDMs**. Reaction time RDM of a representative subject within a learning block (left) and Difficulty based RDM (right).

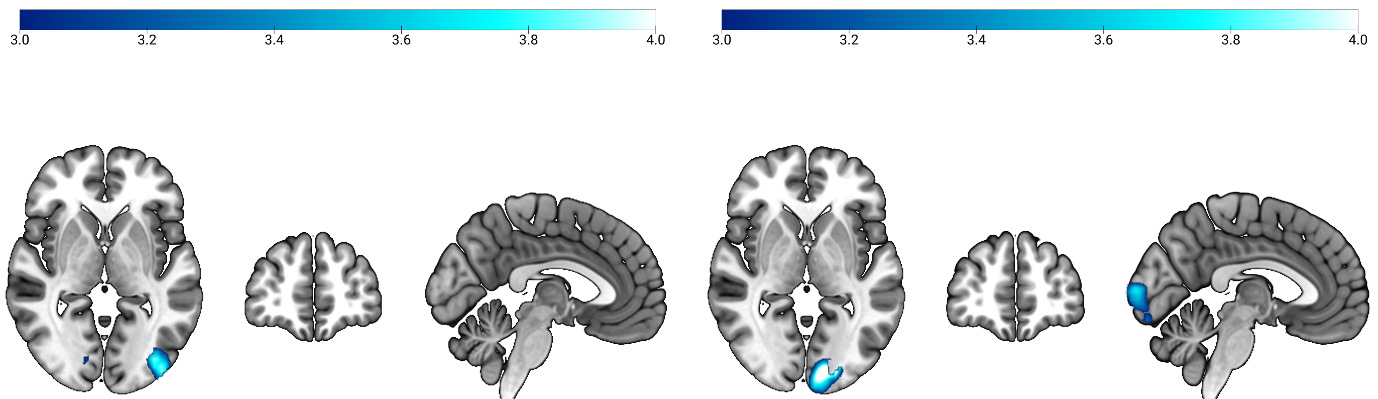

**Supplementary Fig 2: Alternative RDMs Spatial Domain**. Reaction time RDMs (left) and Difficulty based RDMs (right) showing no prefrontal effects.

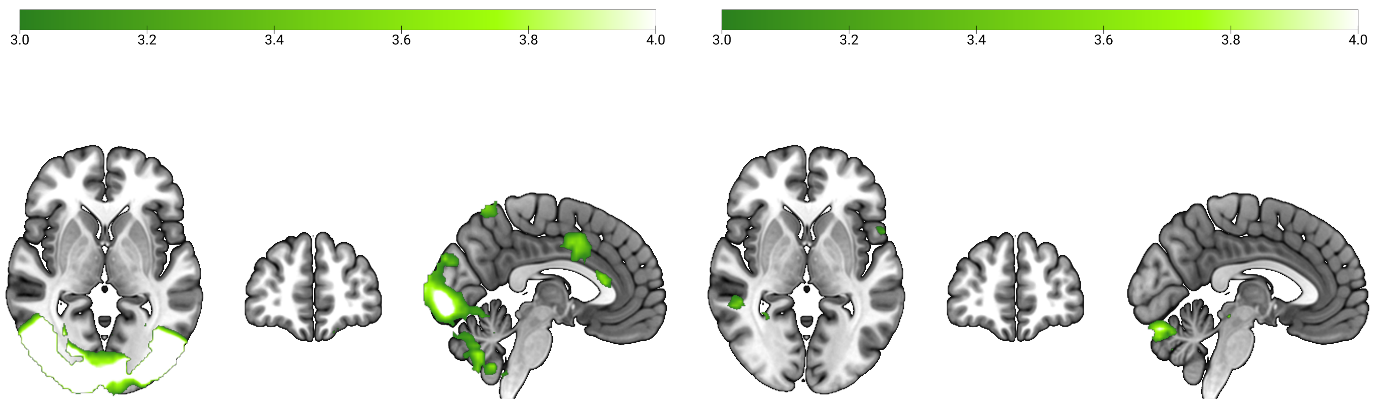

**Supplementary Fig 3: Alternative RDMs Social Domain**. Reaction time RDMs (left) and Difficulty based RDMs (right) showing no prefrontal effects.

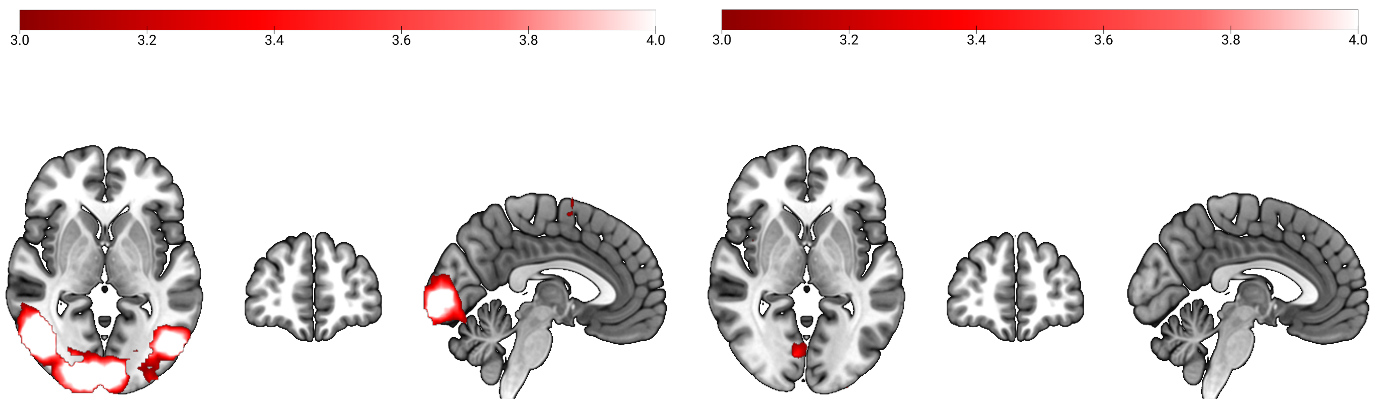

**Supplementary Fig 4: Alternative RDMs Sequential Domain**. Reaction time RDMs (left) and Difficulty based RDMs (right) showing no prefrontal effects.

### Response Times during Learning

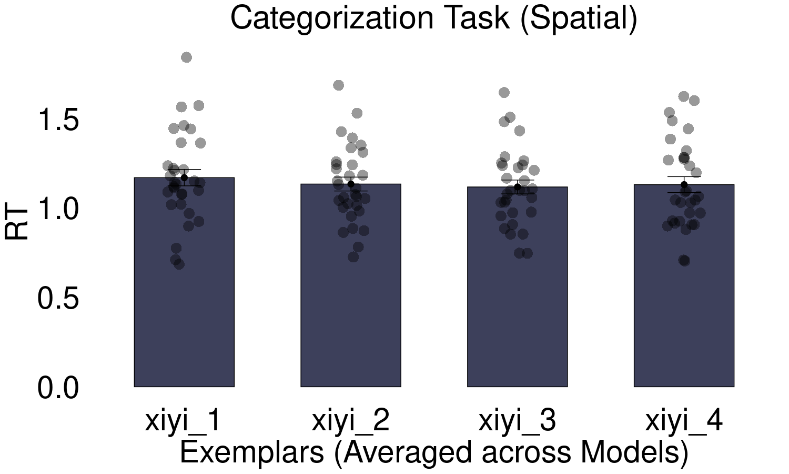

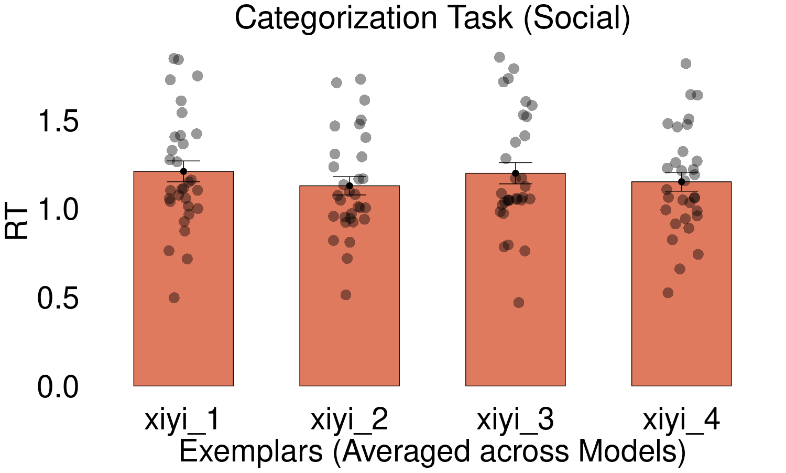

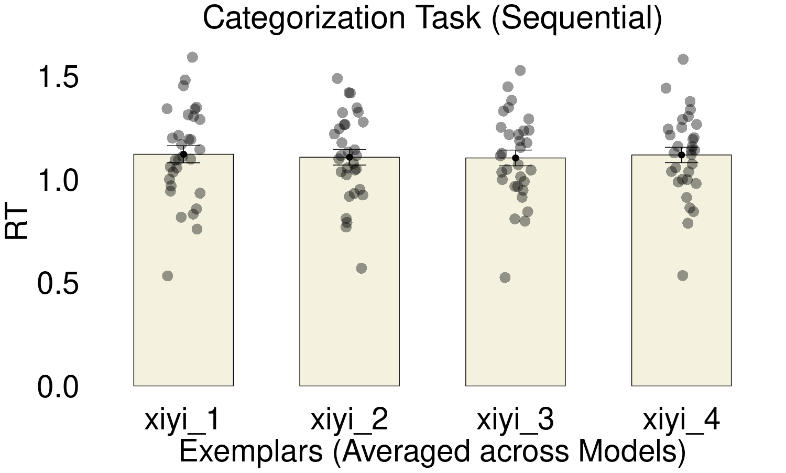

**Supplementary Fig 5: Response times during Learning across domains**.

### Estimated Tolerance used by participants during inference

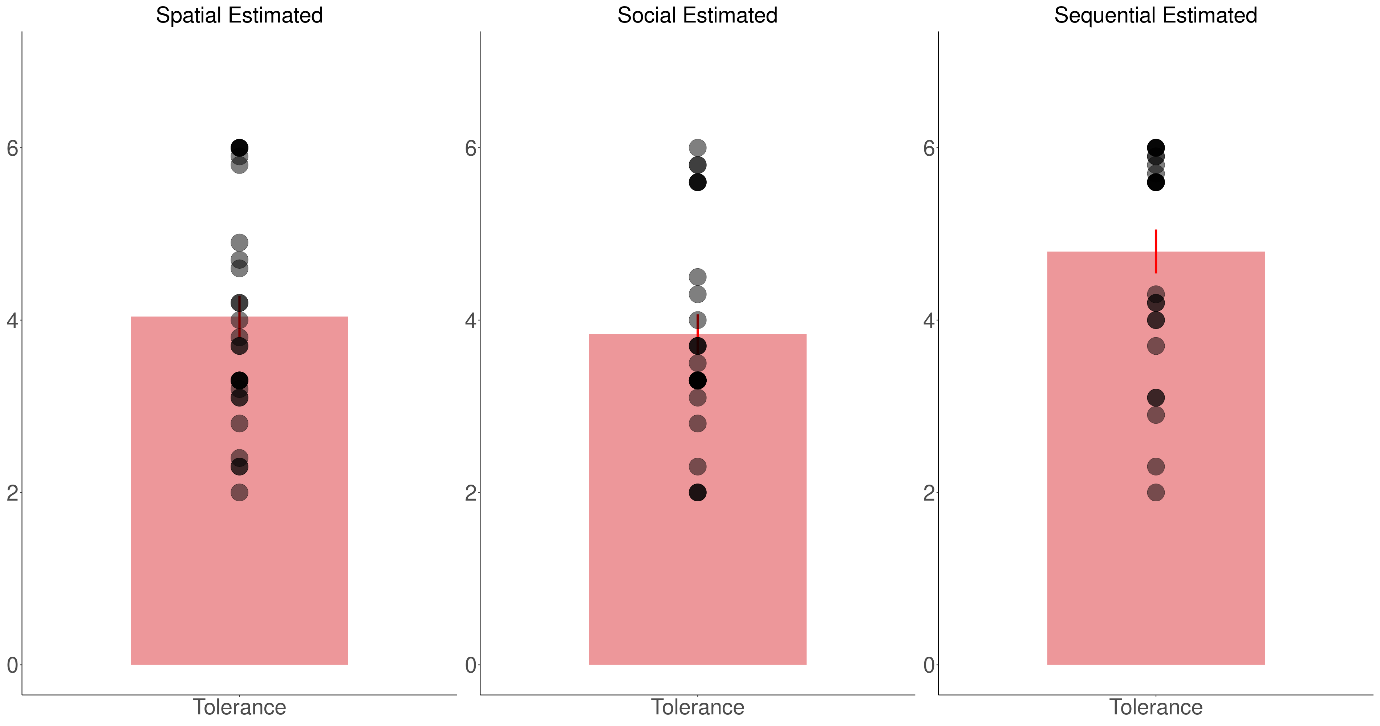

**Supplementary Fig 6: Tolerance used by participants estimated from fitting Monte Carlo model to their full learning and test data in each domain. A higher tolerance would mean more simulated data would be used for making inference after comparison to observed data (under a distance, see Methods). Sequential domain had a higher estimated tolerance across participants than other two domains (see Main Text).**

### Domain-Specific coding in Anterior Temporal Lobe

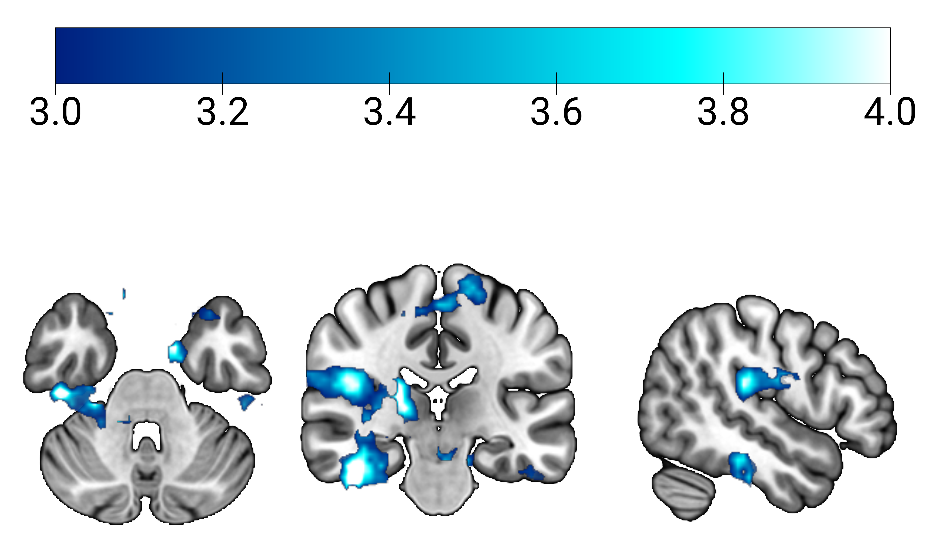

**Supplementary Fig 7: Posterior change RDM (**$\boldsymbol{\delta}_{\boldsymbol{t}}\boldsymbol{)}$ **showing Anterior Temporal Lobe in Spatial domain. Maps are visualized at TFCE corrected p<0.01.**

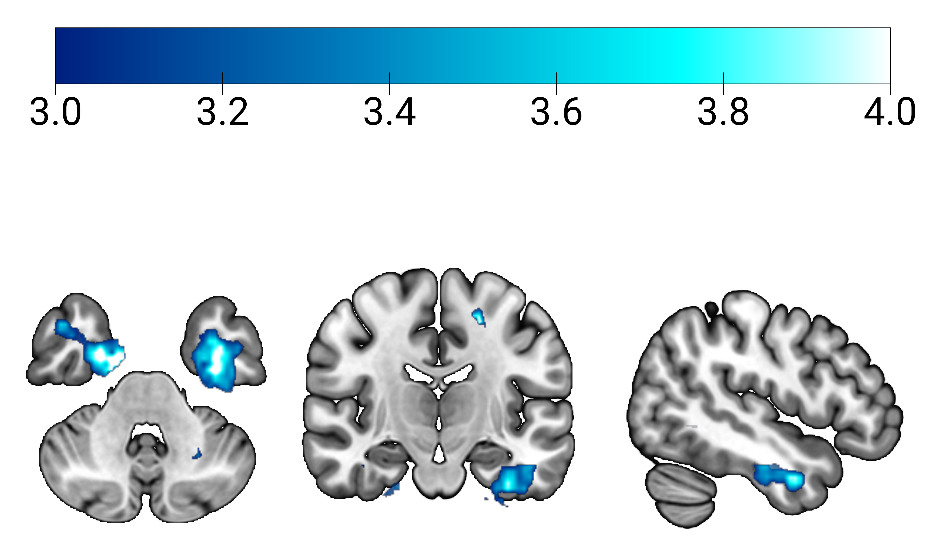

**Supplementary Fig 8: Posterior change RDM (**$\boldsymbol{\delta}_{\boldsymbol{t}}\boldsymbol{)}$ **showing Anterior Temporal Lobe in Social domain. Maps are visualized at TFCE corrected p<0.01.**

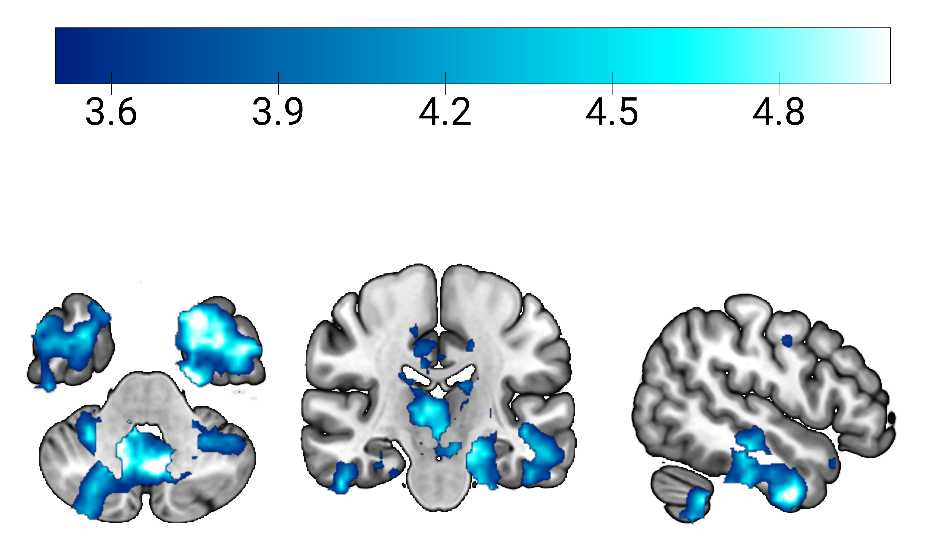

**Supplementary Fig 9: Posterior change RDM (**$\boldsymbol{\delta}_{\boldsymbol{t}}\boldsymbol{)}$ **showing Anterior Temporal Lobe in Sequential domain. Maps are visualized at TFCE corrected p<0.01.**

### Correlations between Process RDMs Across Domains

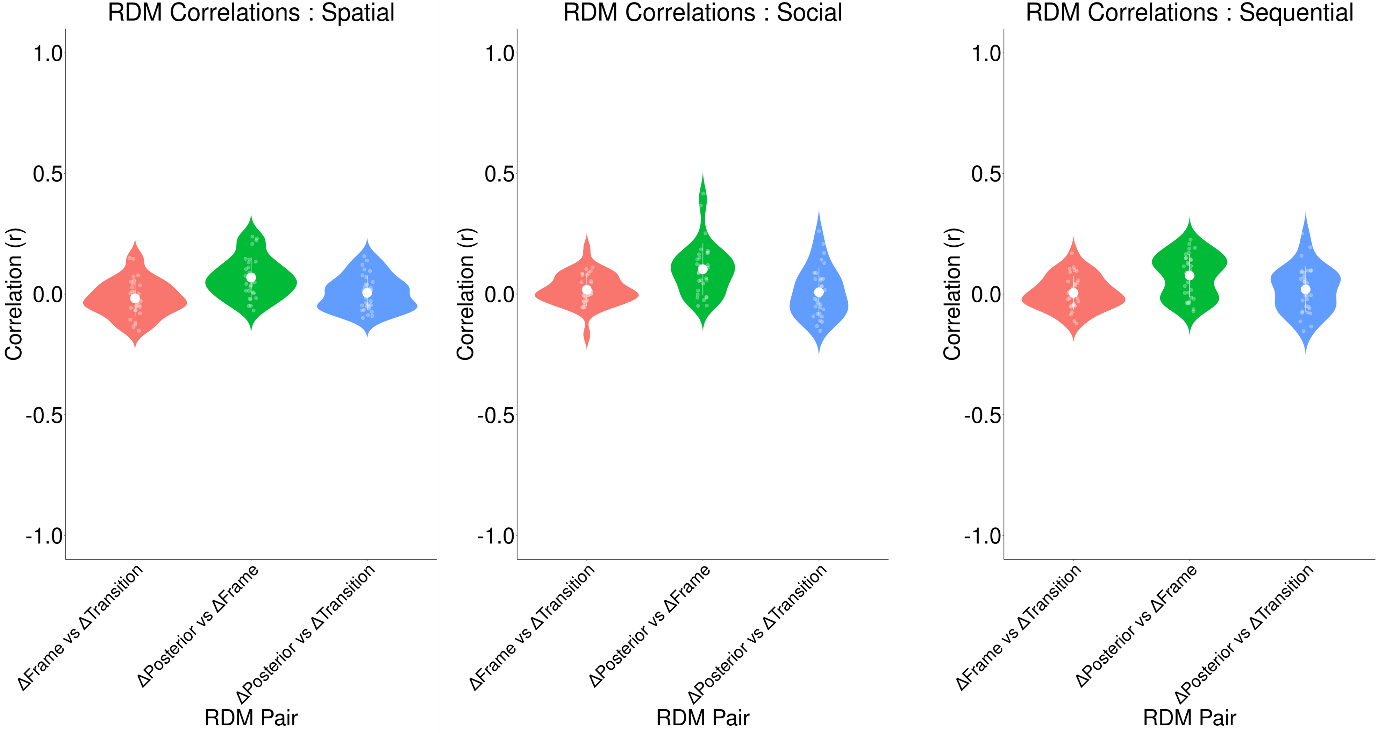

**Supplementary Fig 10: Process RDM correlations in Spatial, Social and Sequential domains**. No significant relationships were found for the three computations despite deriving from the same trial order, suggesting distinct functional profiles. See main text for statistics.
